# Diet quality alters the consequences of gradual phenotypic plasticity for individual performance

**DOI:** 10.1101/2023.03.08.531696

**Authors:** Marine Van Baelen, Alexandre Bec, Erik Sperfeld, Nathan Frizot, Apostolos-Manuel Koussoroplis

## Abstract

Organisms exhibit reversible physiological adjustments as a response to rapidly changing environments. Yet, such plasticity of the phenotype is gradual and may lag behind the environmental fluctuations, thereby affecting the long-term average performance of the organisms. By supplying energy and essential compounds for optimal tissue building, food determines the range of possible phenotypic changes and potentially the rate at which they occur. Here, we assess how differences in the dietary supply of essential lipids modulate the phenotypic plasticity of an ectotherm facing thermal fluctuations. We use three phytoplankton strains to create a gradient of polyunsaturated fatty acids and sterols supply for *Daphnia magna* under constant and fluctuating temperatures. We used three different fluctuation periodicities to unravel the temporal dynamics of gradual plasticity and its long-term consequences on *D. magna* performance measured as juvenile somatic growth rate. In agreement with gradual plasticity theory, we show that in *D.magna*, fluctuation periodicity determines the differential between observed growth rates and those expected from constant conditions. Most importantly, we show that diet modulates both the size and the direction of the growth rate differential. Overall, we demonstrate that the nutritional context is essential for predicting ectotherm consumers’ performance in fluctuating thermal environments.

## Introduction

Understanding how biological processes unfold in fluctuating environments is becoming a central goal in ecological research. This objective is all more urgent in the face of the ongoing climatic change where variability patterns of major environmental factors such as temperature are being profoundly altered. A ubiquitous process by which organisms react to changing environments is phenotypic plasticity. This term refers to the ability of a single genotype to produce alternate behavioral and physiological phenotypes in response to the environmental context. Phenotypic plasticity can be expressed within a generation, allowing organisms to acclimate to their new environment more rapidly than through genetic evolution. However, such plasticity is gradual and takes time to unfold. Recent evidence showed that the trait changes induced by gradual phenotypic plasticity might be slow enough to lag behind the environmental changes that drive it (1–3). Depending on the context and the plasticity mechanism involved, such lag might either penalize (1,3) or promote (4,5) organismal performance relative to expectations in fluctuating environments.

When ectotherms face fluctuating temperatures within their temperature range, a major process involved in phenotypic plasticity is homeoviscous adaptation (6). This physiological mechanism enables the maintenance of an optimal state of cell membrane across the environmental range of temperatures by adjustments of the cell membrane lipid composition. These adjustments compensate for the rigidifying or fluidifying effects on cell membranes of low and high temperatures respectively (6). For most ectotherms, those involve among others, modifications in highly unsaturated fatty acids (HUFA) and cholesterol contents of the cell membrane. HUFA play an important role in acclimation to low temperatures due to their fluidifying effect on the phospholipid bilayer of cell membranes (6). Cholesterol plays an important role at high temperature acclimation, because it rigidifies cell membranes (7,8). Organisms facing warmer environments typically exhibit higher cholesterol concentrations in their tissues (9,10), whereas at lower temperatures they show higher HUFA concentrations (11,12). Cholesterol and HUFA are considered as *essential* compounds for arthropods, that is, they (or their precursors: poly-unsaturated fatty acids [PUFA] and phytosterols, respectively) cannot be efficiently synthesized *de novo* and they must be provided by diet. Yet, dietary supply of these compounds can be limiting in nature, especially for herbivores, thereby constraining the capacity of their cell membranes to acclimate. Under such circumstances their temperature tolerance range may narrow and heat stress may occur at lower temperatures, leading to an earlier onset of a stress response via the production of heat shock proteins (HSP) (13). This type of plastic response promotes survival, but continued overexpression of HSP comes at the cost of decreased growth (5).

Here, we test whether diet quality -as defined by its supply in HUFA (or PUFA precursors) and sterols- can modulate the consequences of *gradual plasticity* of a model ectotherm, the water flea *Daphnia magna*, when facing temperature fluctuations. As we explain below (see *Theory* section), we hypothesize that HUFA- and sterol-rich diets should promote membrane plasticity. On one hand, growth under fast fluctuating temperatures should be lower than the average growth expected from fully acclimated phenotypes. On the other hand, diets lacking these compounds should constrain membrane plasticity, but elicit heat stress responses when temperatures exceed the optimal temperature for growth. In this case, growth under fast fluctuating temperatures should be higher than the average growth expected from fully acclimated phenotypes. Below, we explain how we used the theoretical framework of *gradual plasticity* (1,3) and the concept of time-dependent effects (5,14) to construct our experimental design and test our hypotheses.

### Theory

The gradual plasticity framework (*see* 1 for a detailed presentation)) is based on the concept of the performance landscape (Fig. 1A) which describes how an organismal performance trait (e. g. growth rate), *P*, depends on the current environment (here temperature, *T*) and its current phenotype, Φ, so that *P* = *f*(*T*, Φ). The diagonal line along a performance landscape indicates a phenotype matching its temperature (Φ) or in other words, an organism experiencing a given temperature for a time sufficiently long to allow its acclimated phenotype to be fully expressed. All surface above or below that diagonal indicates a mismatch between the phenotype and the current environment (Φ ≠ Φ*). An acute change in temperature can be visualized as a deviation from the diagonal line parallel to the temperature axis. The process of phenotypic plasticity can be viewed as a parallel movement along the Φ axis and towards the diagonal line either increasing performance over time (Fig. 1B blue line) or decreasing it (Fig. 1B red line) two phenomena termed *beneficial* and *detrimental* plasticity, respectively (*sensu* 3).

**Figure 1:**
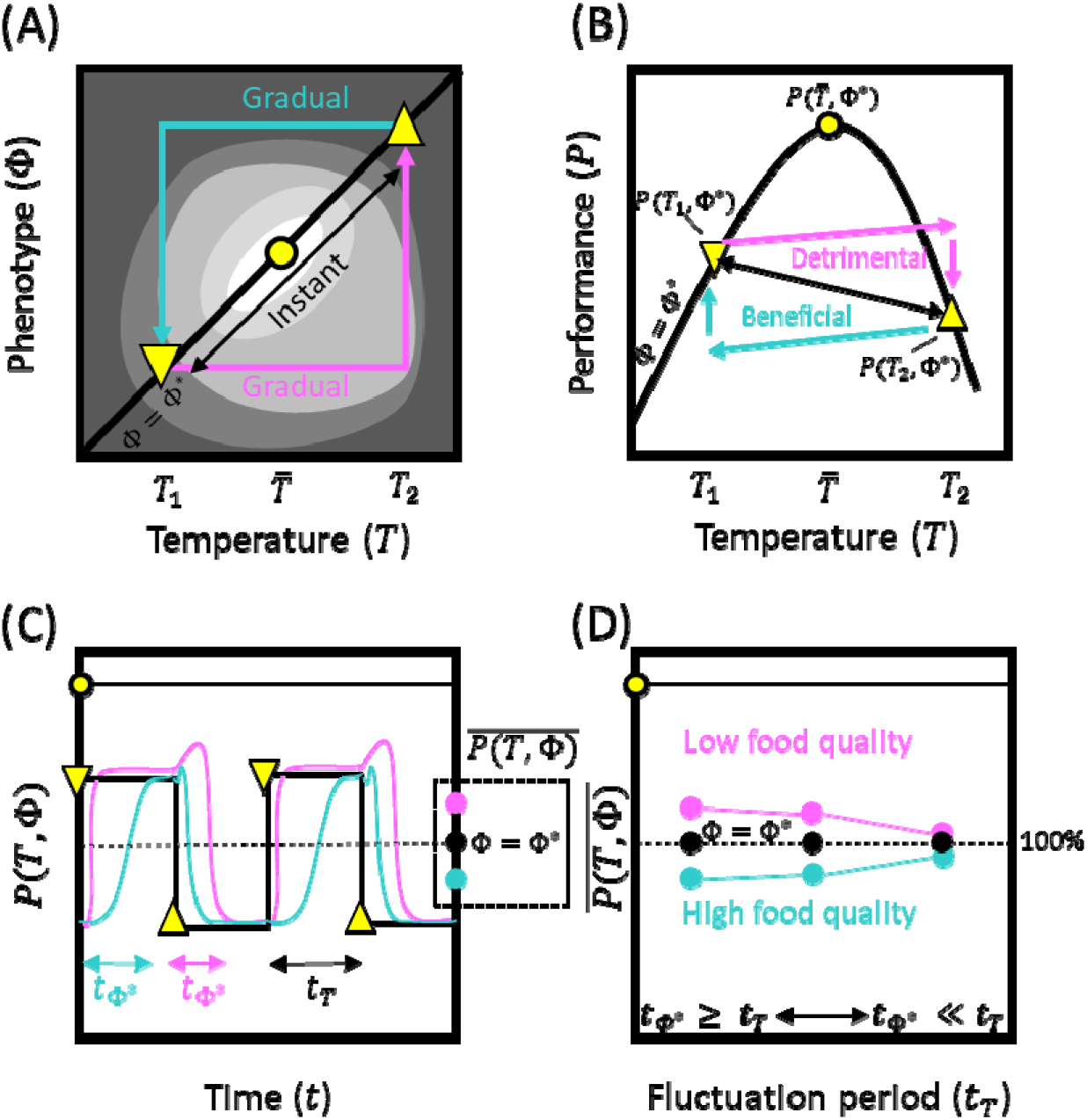
(adapted from Fey *et al.* 2021) **Performance landscapes** (**A**) describe how organismal performance (contours) depends on its current phenotype (Φ) and temperature (*T*). The diagonal line is the acclimated **thermal performance curve** (TPC) (**B**). When changes are gradual, an individual may spend significant time away from the acclimated phenotype (Φ*) with consequences on **time-averaged performance**, 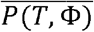 (**C**). When detrimental [beneficial] plasticity dominates, the time-averaged performance should be higher [lower] than 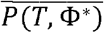. The differences depend on how fast the acclimated phenotype is reached (*t*_φ*_) relative to the time scale at which the temperature fluctuates (*t_T_*) leading to **time-dependent effects** (**D**). Note that beneficial and detrimental plasticity in (A) and (B) are displayed together for convenience, they do not necessarily co-occur. Also, in (C) and (D), the axis is normalized with regard to 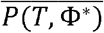. In absolute terms, a high-quality diet yields higher performance than a low quality one.

While the performance landscape is a very useful concept, measuring such landscapes directly is a daunting task, except perhaps for organisms with minimal spatial requirements and very short generation times such as microbes (*see* 1,3). Yet, phenotypic plasticity can also be detected indirectly by quantifying the deviation between experimental observations of time-averaged performance in a fluctuating environment, 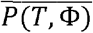, and the predicted time-averaged performance under the assumption that phenotypic plasticity is instantaneous or nonexistent i.e. 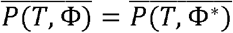. To achieve that, one needs to measure *P*(*T*, Φ*) across the range of temperatures of interest. The resulting curve is the acclimated temperature performance curve (TPC) (Fig. 1B) and corresponds to a slice along the diagonal line of the performance landscape (Fig. 1A). When phenotypic plasticity is gradual, the phenotype might lag behind the environmental fluctuations and Φ might never reach Φ* or reach it for a negligible relative amount of the total time considered (Fig. 1C). In the case of beneficial phenotypic plasticity, this implies that for most of the time over which performance is averaged (Fig. 1C; blue line), the organism under-performs relative to the fully acclimated phenotype hence yielding 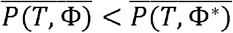. In the case of detrimental plasticity, the organism might never experience the complete decrease in performance caused by a prolonged exposure to a given temperature (Fig. 1C; red line). In this case, for most of the time over which performance is averaged the organism over-performs relative to the fully acclimated phenotype hence resulting in 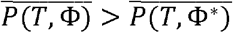. However, if the environmental fluctuations are slow enough relative to the time required for Φ to reach Φ*, the relative amount of time spent away from Φ* becomes negligible and 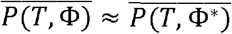 (Fig. 1 D). Hence, the value of 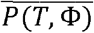 and its difference from 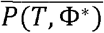 depends on the time scale at which temperature fluctuates, a phenomenon known as *time-dependent effect* (5,14–16). For the reasons exposed above, we hypothesized that high quality diets (with regard to essential lipid supply) should promote beneficial phenotypic plasticity when facing temperature fluctuations. On the other hand, low quality should yield detrimental phenotypic plasticity. If our hypotheses are correct: when facing periodic temperature fluctuations, (*1*) 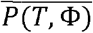 of consumers fed with high quality diets should be lower than their 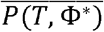, (2) 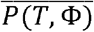 of consumers fed with low quality diets should be higher than their 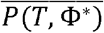, (*3*) for both diets, 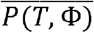 should converge towards 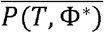 for longer temperature fluctuation periods (Fig. 1D). While not being a definitive proof that the mechanisms invoked here (homeoviscous adaptation, HSP) are in action, confirming our hypotheses would indicate that diet can modulate the consequences of phenotypic plasticity for ectotherms facing temperature fluctuations.

## Methods

### Experimental organisms

We used a clonal line of the freshwater cladoceran *Daphnia magna* (IRTA-6, hereafter *Daphnia*) as our experimental model organism. The stock cultures of *Daphnia* were kept in Volvic water© at 20°C on a 12:12 day/night cycle and fed with the green algae *Chlamydomonas reinhardtii* at 3 mg C L^-1^ (well above the limiting level that is reported at 0.7 mg C L^-1^(17).

We used three phytoplankton species to create a gradient of PUFA, HUFA and sterol supply for exploring how dietary constraints affect thermal performance. *Cryptomonas sp.* (strain SAG 2680), a cryptophyte rich in HUFA (especially eicosapentaenoic acid, EPA, 20:5ω3) and sterols (18,19), constitutes an excellent quality food for *Daphnia* (20). The chlorophyte *Chlamydomonas reinhardtii* (SAG 7781) has lower sterol concentrations than *Cryptomonas* (19), lacks EPA, but contains its precursor alpha linoleic acid (ALA; 18:3ω3) (21). *Chlamydomonas* is considered here as an intermediate quality food (20). *Synechococcus sp.*, a cyanobacterium, lacks both PUFA and sterols and is therefore a poor quality food for *Daphnia* (20). As *Synechococcus sp.* alone does not allow *D. magna* growth, we supplemented it with 10% of *Chlamydomonas*. All phytoplankton species were grown on sterilized WC medium (22) with vitamins at 20°C under permanent light with a dilution rate of 0.25 per day in aerated 5L vessels.

### Fluctuating temperature experiments

*Daphnia* were fed on three different diets and incubated in climate chambers (POL-EKO-APARATURA SP.J, ST3/3C SMART) with temperatures alternating between 20°C and 28°C with 3 different periodicities while trying to keep the same mean temperature and variance across treatments (Fig. 2B-D). The growth experiments lasted 144 hours (6 days) in total. In the first periodicity treatment (F12) the temperature was shifted from 20°C to 28°C approximatively every 12h. In the second treatment, five 24h periods of alternating 20°C or 28°C where followed by a last 18h period at 28°C (average fluctuation period 22h, named F24 for simplicity). The last treatment (F48) was composed of three periods of approximatively 48h alternating between 20°C and 28°C. We used HOBO© temperature loggers (MX2200, Onset) immerged in experimental vessels filled with water in order to measure the actual temperature experienced by *Daphnia* during the experiment. The three periodicity treatments yielded similar mean temperatures, but somewhat different variances (22.67-23.53°C and 10.1-12.2°C^2^, respectively, see Sup. data Table S1).

**Figure 2:**
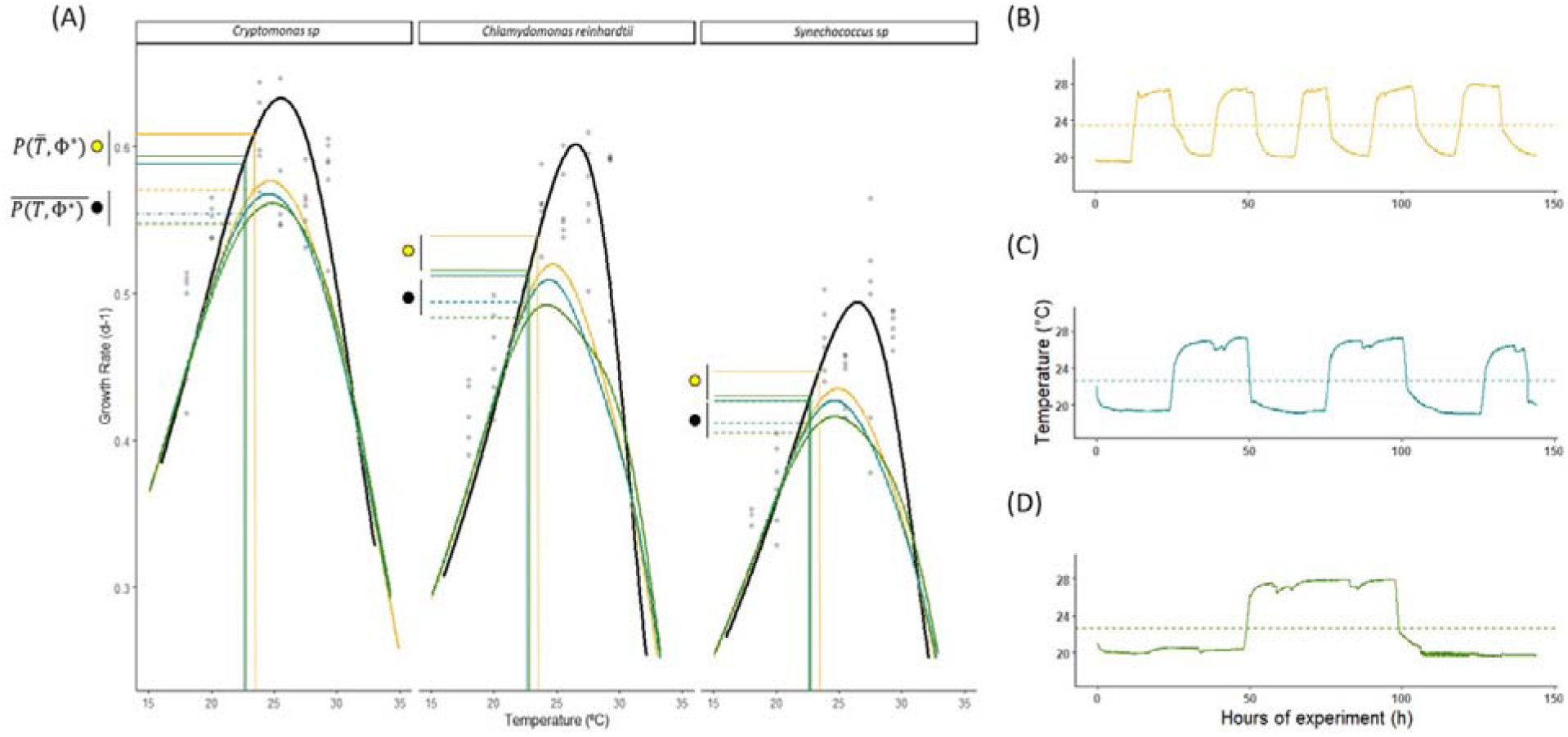
(A) **Thermal Performance Curves** TPC (in black) for *Daphnia magna* growth rate (d-1) submitted to different food treatments: Cryptomomas sp., Chlamydomonas reinhardtii and Synechococcus sp. on constant temperature. **Theoretical time-averaged performance** 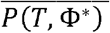 on the different temperatures for the 12’hours (gold), 24’hours (blue), 48’hours (green) treatments. The **Theoretical time-averaged performance** 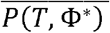 and the **mean temperature performance** 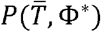 for the mean temperature of each treatments are indicated with respectively dotted and full lines. The differences in these lines emerged due to the slight differences in mean and variance in the different fluctuation regimes (*see sup data* Table. 1) (B-D) Water temperatures achieved in (a) 12’hours (b) 24’hours (c) 48’hours thermal variation treatments, where the full lines indicate experience temperature from 20°C to 28°C and the dotted line represents the mean temperature of the treatment. Lines are temperatures taken every minute ± 0.1°C from Hobo.

The experiments were initiated with third clutch synchronous (< 6h old) *Daphnia* neonates from the same clonal line born from mothers acclimated at 24°C and fed with *Chlamydomonas.* The neonates were randomly collected and fed *Chlamydomonas* for 5 hours. After the 5h of pre-feeding, they were distributed to 4 replicate 600ml vessels of 15 individuals each (4 replicates x 3 diets x 3 fluctuation periods = 48 vessels), and an additional three replicates of 30 organisms were used for initial dry weight determination (W0). In each of the 600ml vessels, organisms were raised in Volvic© water renewed every day with a daily food supply of 3 mg C L^-1^ during the 6 days of the experiment. At day 6 (144hours), 12 individuals per replicate were randomly chosen and placed individually into pre-weighed aluminum containers, dried for 48h at 60°C and weighed with a precision of 0.001mg using a microbalance (Sartorius; ME36S; Goettingen-Germany) to determine their dry weight. The somatic growth rate was calculated as:

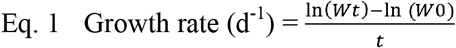

Where W0 is the average individual weight (μg) of neonates at time 0h and Wt the average individual weight (μg) after 6 days *t* (days).

### Predicting performance in fluctuating environments

In order to have a reference for performance without temperature fluctuations and to model the Thermal Performance Curve (TPC), subsequently used to predict the effect of temperature variance on *Daphnia* performance (*see* Fig. 1), we additionally grew *Daphnia* at constant temperatures of 18°C, 20°C, 24°C, 26°C, 28°C and 30°C, fed the three diets of different quality described above. For logistical reasons, growth experiments were performed independently for each temperature using the same clonal line and a strictly identical protocol. The experiments were initiated with third clutch neonates from mothers acclimated at each experimental temperature and fed with *Chlamydomonas.* Neonates were randomly distributed to 6 replicate vessels containing each 15 individuals for each diet treatment (6 replicates of 15 individuals x 3 diets x 6 temperatures) and three additional replicates of 30 organisms were used for initial dry weight determination (W0). The organisms were maintained in Volvic© water renewed each day with a daily food supply of 3 mg C L^-1^ for 6 days. Every day, 6 individuals were randomly chosen, placed individually into pre-weighed aluminum containers, dried for 48h at 60°C and weighed as described in the previous section to calculate the somatic growth rate from day 0 to day 6.

### Data analysis

We used general linear models to test whether the diet and temperature fluctuation periodicity affect somatic growth rate. The variance explained by each factor and their interaction was estimated using a two-way ANOVA (type II) followed by a Tukey post-hoc test. The normality and homoscedasticity of the residuals were verified using the Shapiro-Wilk and Levene tests, respectively. In order to calculate the theoretical time-averaged performance 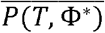 we first obtained the constant temperature condition TPC, 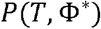, by comparing different models (*see* sup. data Fig.S1-S2) and by fitting the Sharpe-Schoolfield model to the growth rate data obtained for each diet treatment (Fig.2A) using:

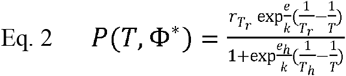

Where *T* is the body temperature (K); *r_T_r__* is the growth rate at the reference temperature *T_r_*= 288°K (15°C), *e* the activation energy (eV), *e_h_* the de-activation energy (eV) at high temperature (K) *T_h_*, and *K* is the Boltzmann’s constant (8.62e-05 eV K^-1^). The Sharpe-Schoolfield model was fitted using R V 4.0.3 (R Core Team 2020) and the rTPC package (23,24). The model equation explains the thermal sensitivity of a rate by enzymes deactivated at both extreme-high and low temperature (23,24). However, here, we only used the high temperature enzyme inactivation, because low temperature required multiple rate measurements at low temperatures for inferring accurate parameter estimates (23). The theoretical time-averaged performance 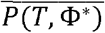 for each diet and temperature fluctuation periodicity combination can be calculated using the numerical integration of equation 2 (25):

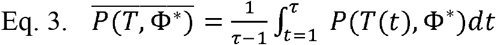

where *T*(*t*) is the temperature time-series obtained in the three periodicity treatments (with 1-minute resolution; see figure 2A-D) and *τ* the number of minutes per time series. The means and associated 95% confidence intervals (C.I.) of *P*(*T*, Φ*) and 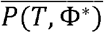 were estimated by non-parametric bootstrapping (n=1000) of the constant temperature growth rate datasets (26). The significance of the differences between the observed performances under fluctuating temperature condition, 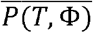, those predicted by time-averaging, 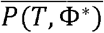, and those expected in a constant environment with the same mean temperature 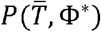 was tested using one sample T-test bootstrapping.

## Results

### Effect of thermal fluctuations

Relative to constant temperature, thermal fluctuations have a significant effect on growth responses depending on the diet and the periodicity treatment (Fig. 3). On *Cryptomonas*, growth rates 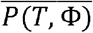 are 9.70% to 12.63% lower than performances on constant mean temperature, 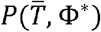 (Sup. data Table S2, Student test: t_10_=13.68, p<0.001) and, similarly, on *Chlamydomonas*, growth rates under thermal fluctuations are 5.25% to 10.29% lower than 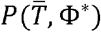 (Sup. data Table S2, Student test: t_11_=4.32, p<0.005). On *Synechococcus*, only temperature fluctuations every 12 hours caused a growth decrease of 2.89% relative to the mean constant temperature treatment performance (Sup. data Table S2, Student test: t_10_=2.36, p<0.05).

**Figure 3:**
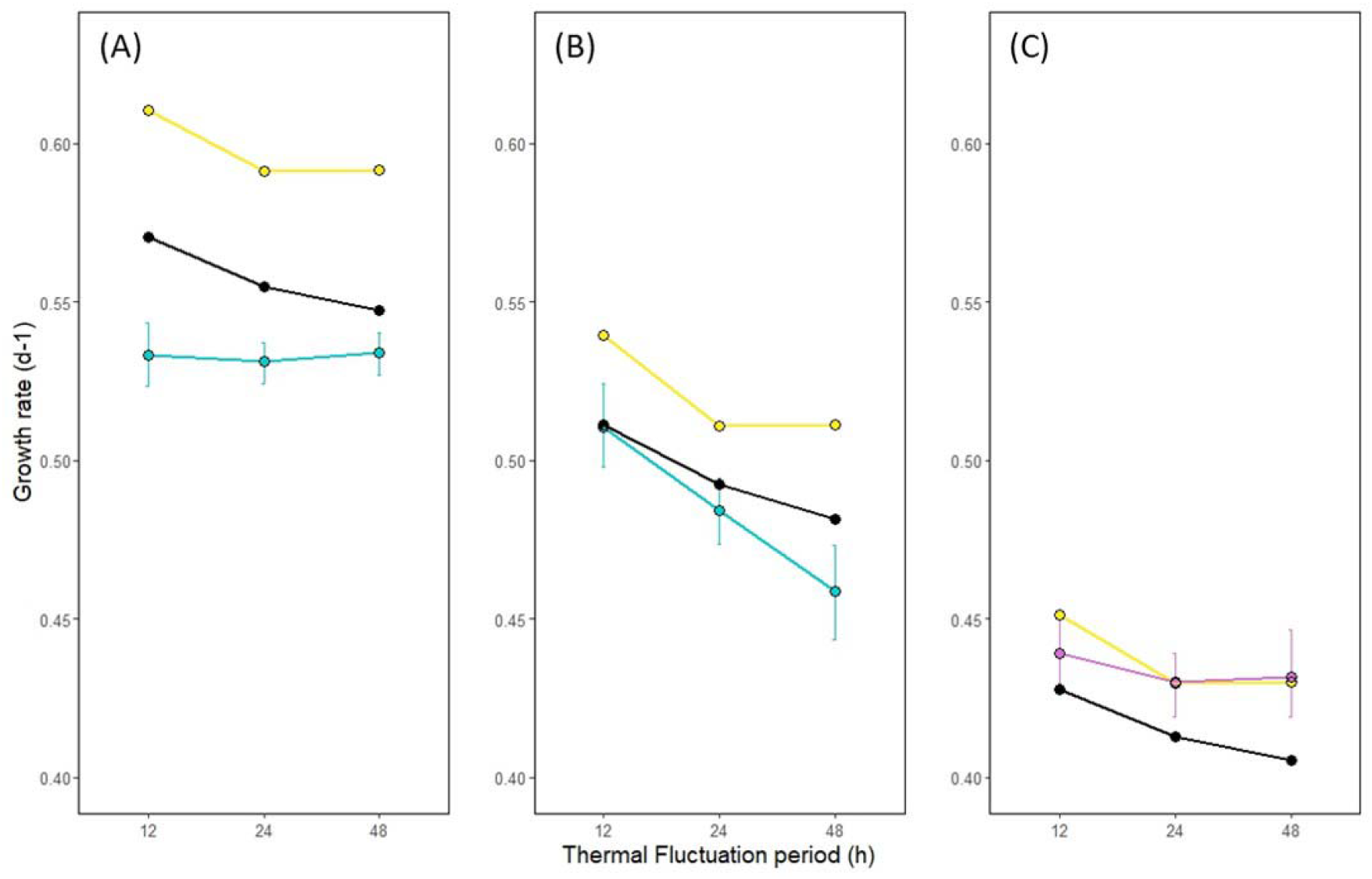
Daphnia magna growth rate (day^-1^) under different diets ((A) *Cryptomonas sp.*, (B) *Chlamydomonas reinhardtii.*, (C) *Synechococcus sp.)* for each thermal period treatment (12’hours (*12*), 24’hours (*24)* and 48’hours treatments (*48*)). **Observed growth rate** with under acclimated plasticity (blue) and over-acclimated plasticity (pink), Theoretical **time-averaged performance** (black) and the **mean temperature performance** (yellow).

### Observed and predicted growth

In most of the thermal periodicity treatments, the observed performances under fluctuating temperature conditions, 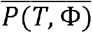, differ from those predicted by time-averaging performance, 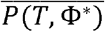 (Fig. 3). On the *Cryptomonas* diet, the observed growth rate of *Daphnia* is significantly lower than the predicted performance (Table 1, Student test: t_10_=6.64, p<0.001). The observed growth on *Chlamydomonas* is significantly lower (4.8%) than the predicted growth only for the 48h periodicity (Table 1, Student test: t_11_=2.96, p<0.005). For *Daphnia* grown on *Synechococcus*, 24h and 48h fluctuation treatments are significantly higher than the predicted performances (Table 1, Student test: t_11_=3.57, p<0.001).

**Table 1:**
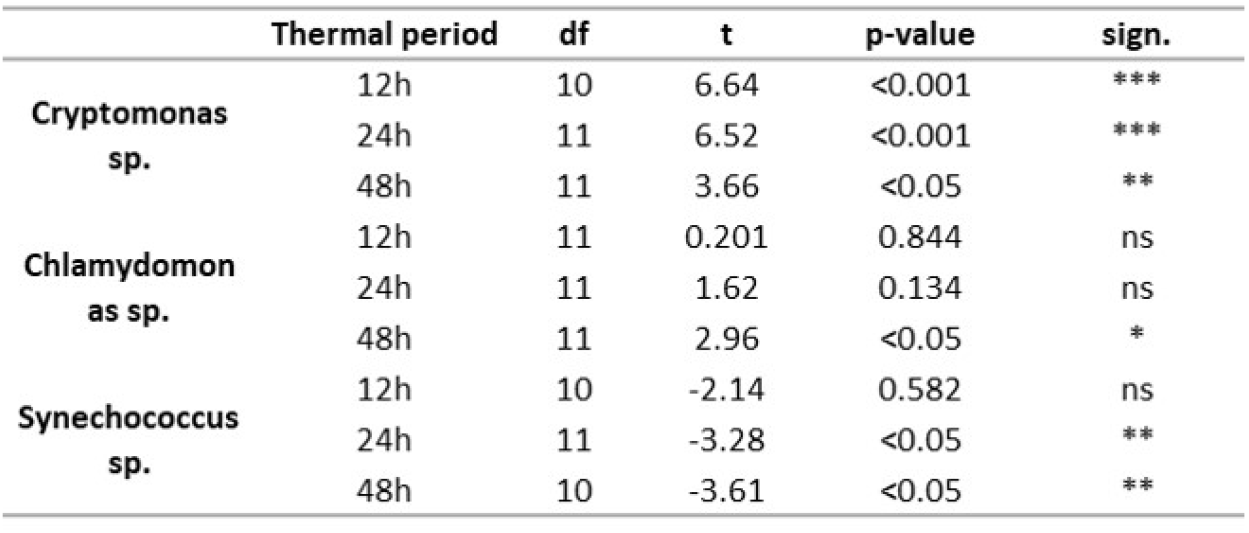
Student test bootstrapping. Observed growth rate between theoretical time-averaged performance results of Daphnia magna growth rate (day^-1^) under different diets (Cryptomonas sp., Chlamydomonas reinhardtii., Synechococcus sp.) for each periodicity treatment. Thermal fluctuation periodicity of 12’hours (***12h***), thermal fluctuation periodicity of 24’hours (***24h***), thermal fluctuation periodicity of 48’hours (***48h***).

### Food and periodicity effect

There was a significant thermal regime by diet interaction (ANOVA: F_4,96_= 5.9, p < 0.001) on the growth rate of *Daphnia* (Table 2). In all treatments, the highest growth rates were observed on the *Cryptomonas* diet, followed by the *Chlamydomonas* and *Synechococcus* diet. Periodicity had no effect on growth rate for *Cryptomonas*, but performances seem to converge towards the time-averaged growth, 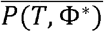 (Fig. 3). For *Chlamydomonas*, observed growth rates significantly diverge from 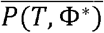 with increasing thermal periodicity (Fig. 3 and sup. data Table S3, post-hoc ANOVA slow/fast period; F_96_=6.32, p<0.001). However, on *Synechococcus* sp., even if there is no significant effect of the periodicity, observed growth rates seem to diverge from 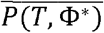 (Fig. 3).

**Table 2:** ANOVA results (Type II tests) of Daphnia magna growth rate (day^-1^) under different diets (Cryptomonas sp., Chlamydomonas reinhardtii., Synechococcus sp.) for each periodicity treatment. The ANOVA test describes diet quality and periodicity treatment effect but also the interaction between the two on D. magna growth rate (day^-1^).

|  | SS | df | F | p-value |
| --- | --- | --- | --- | --- |
| **(Intercept)** | 24.5184 | 1 | 61524 | <0.001 |
| **Periodicity** | 0.0066 | 2 | 8.34 | <0.001 |
| **Diet Quality** | 0.1701 | 2 | 213.39 | <0.001 |
| **Diet Quality \* Periodicity** | 0.0094 | 4 | 5.9 | <0.001 |
| **Residual Error** | 0.0383 | 96 |  |  |

## Discussion

Phenotypic plasticity can be detected from deviations between experimental observations of performance in a fluctuating environment, 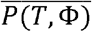, and the predicted time-averaged performance, 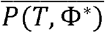. Here, we find growth rates lower or higher than expected from the acclimated thermal performance curve (*TPC*) depending on diet quality. This mismatch between the observed and the predicted performances points towards the capacity of *Daphnia* to express *gradual plasticity* (27) and that the current phenotype (Φ) lags behind the tested temperature fluctuations (1,3). Furthermore, our results suggest that the difference between 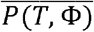 and 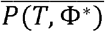 might depend on the time scale at which temperature fluctuates, a phenomenon known as *time-dependent effect* (5,14,16).

Our experimental results support our hypothesis that diet quality can modulate the consequences of *phenotypic plasticity* on *Daphnia* somatic growth rate: On the high-quality food *Cryptomonas*, our results show that the observed growth 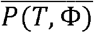 is consistently lower than the predicted time-averaged performance, 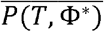. On the other hand, for the low-quality food *Synechococcus*, 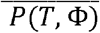 was equal (12h) or higher (24h and 48h periodicities) than 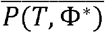. For the intermediate quality food *Chlamydomonas*, 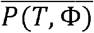 is equal (12h and 24h) or lower (48h) than 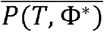. We argue that this pattern can at least be partially explained by the contents of sterols, PUFA and HUFA in the different diets. *Cryptomonas* is rich in these compounds involved in homeoviscous adaptation (18,19) and therefore enables beneficial phenotypic plasticity (*sensu* 1) towards temperature changes. As demonstrated by Fey et al. (2021), such type of plasticity can penalize growth rate relative to the expectations in the fluctuating environment. On the other hand, the lack in these compounds, more particularly sterols in the *Synechococcus* diet, probably constrains the homeoviscous adaptation capacity of *Daphnia* and may in turn increase the susceptibility of the organism to high temperatures and heat stress (10,28).

While no link between sterol limitation and HSP expression was found in sterol auxotroph yeasts (29), in mammalian cells, the hyperfluidization of cell membranes (such as the one that could be induced in our sterol limited daphnids at higher temperatures) lowers the set-point temperature at which HSP synthesis is initiated (30). By analogy to homeoviscous adaptation, HSP expression can also be viewed as gradual phenotypic plasticity which benefits certain traits such as survival to future heat stress, but comes at a cost for other traits such as growth (13,31). The thermal performance curves used to predict 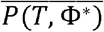 under fluctuating conditions for *Synechococcus* were build using somatic growth measured after six days of exposure to constant temperatures. This time frame is hence encompassing any potential negative effects of HSP overexpression at higher temperatures. Under fluctuating thermal conditions however, the organisms are only partially exposed to stressful higher temperatures, which may limit HSP accumulation and its deleterious effects on growth rate (4,5) and lead to 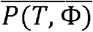 being higher than 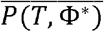.

For the *Cryptomonas* diet, our results also support our hypothesis that observed growth rates 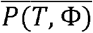 should converge toward the expected time-averaged performances 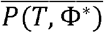 with decreasing fluctuation period. Moving from 12h to 48h periods, the organism experiences constant temperatures over longer duration, thus allowing the organism to fully express and benefit from the acclimated phenotype (Φ*), so that 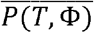 tends towards 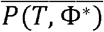. A possible explanation for this pattern could be the HUFA concentration of *Daphnia*’s cell membranes lagging behind in faster fluctuating thermal environments. When available in the diet, *Daphnia* tends to incorporate large amounts of HUFA in their tissues (11,12,32). At cooler temperatures, as at the beginning of the fluctuation experiments, this accumulation of dietary HUFA is beneficial as they contribute to compensate for the rigidifying effect of low temperatures on cell membranes (6). At higher temperature, however, an excessive accumulation of HUFA could have detrimental effects on *Daphnia* performance (33,34). Hence, daphnids should reduce their HUFA excess in their cell membranes through homeoviscous adaptation, a plasticity process that may take some time. In unicellular eukaryotes, remodeling of membrane lipid composition may take between minutes and hours to complete (35). In small invertebrates such as collembola, fatty acid composition continues to change several days after temperature shift (36,37). To our knowledge, the time scale of cell membrane lipid composition remodeling in *Daphnia* is not known, but it likely lies within the daily scale rather than the hourly scale. Hence, daphnids in our fluctuating treatments could have experienced the detrimental effect of HUFA excess, accumulated while feeding on *Cryptomonas* during the initial cooler phase, during the warmer phases. This putative detrimental effect might explain our observations: at shorter periodicities the organisms might have spent a larger portion of time suffering from membrane HUFA excess leading to a larger difference between 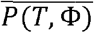 and 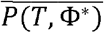. As periodicities become longer, this portion of time decreases, giving the organisms more time to adjust membrane lipid composition and thus leading to a convergence between 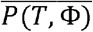 and 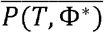.

For the *Chlamydomonas* and *Synechococcus* diets, our hypothesis that 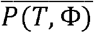 should converge towards 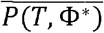 with decreasing fluctuation period was not confirmed as we observed the opposite pattern. This apparent contradiction could be explained by the possibility that the response of 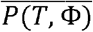 to the fluctuation period (or frequency) of temperature is unimodal over the range of the studied periods (or frequencies). For example, (38) showed that the time-averaged growth rate of photosynthetic microorganisms exposed to fluctuating light intensity is a unimodal function of light fluctuation frequency. In their experiment, the maximal difference between observed and predicted growth rate was recorded at intermediate light fluctuation frequencies. Both high and low frequencies led to a convergence of the two values. The authors proposed an explanation based on concurrent physiological mechanisms (Calvin cycle and RuBisCo kinetics) operating at different time scales. Identifying similar processes here is beyond the scope of the study but warrants further research. If the time-averaged performance of our daphnids is a unimodal function of temperature fluctuation period, as in the preceding example, our results could be explained by diet quality changing the shape of the unimodal response. As a consequence, the temperature fluctuation periods explored here for the *Chlamydomonas* and *Synechococcus* diets might lie within the decreasing part of the function (i.e. the range of fluctuation periods where 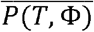 correlates negatively with fluctuation period), while lying in the increasing part for the *Cryptomonas* diet.

## Conclusion

Understanding and predicting organism performances in variable and changing environments is central to forecasting future ecological outcomes (39). Unravelling the temporal dynamics of phenotypic plasticity and their consequences on species traits appears as an important step towards that goal (1,27). Here, we show that food quality can modulate the temporal dynamics of phenotypic plasticity under environmental fluctuations. We are aware that our results are based on partially arbitrary temperature fluctuation patterns and cannot be directly transferred to nature, although both the periodicities and the amplitude are ecologically relevant. Rather, our results need to be viewed as a proof of concept that diets with substantially varying PUFA and sterol contents elicit different consequences of phenotypic plasticity on the growth rate of ectotherms exposed to thermal fluctuations. At this point, it is also important to note that all the physiological mechanisms discussed above are putative and that the actual mechanisms underlying our results are unknown. Yet, based on current knowledge, the dietary modulation of the capacity for homeoviscous adaptation is a good candidate. Furthermore, this mechanism is not only involved in temperature acclimation but also in acclimation to salinity, pH, hydrostatic pressure as well as hypoxia (40), all of which are environmental factors that can tremendously fluctuate during the individual lifetime of an organism. This broadens greatly the potential importance of lipid nutrition as a modulator of phenotypic plasticity and its consequences on organismal performance. In aquatic environments, essential lipid supply to consumers is driven by microbial communities (phytoplankton, biofilms) and varies greatly in space and time (41). Under the pressure from global changes such as warming and eutrophication, these microbial communities are strongly altered and increasingly dominated by cyanobacteria (42–45) lacking both HUFA and sterols (46) with potentially far reaching consequences on the consumer’s responses to alterations in variability patterns of their physical environment.

## Supporting information

Data and Script

