## Supplementary material for "Diet quality alters the consequences of gradual phenotypic plasticity for individual performance": Data and Script: boatman.R

```
library(boot)
library(car)
library(rTPC)
library(nls.multstart)
library(broom)
library(tidyverse)
library(patchwork)
library(minpack.lm)
library(rstatix)
library(rcompanion)
setwd("C:.../Supplementary_data/data")
data.0<- read.table("growth_rate_constant.txt", sep="\t", dec=",", header=TRUE)

data.0$M<-data.0$M*1000####mass in mg
mi=8.8##############mean initial mass
data.0$rate<-(log(data.0$M)-log(mi))/(data.0$time*0.0416667)

label_facets_num <- function(string){
  len <- length(string)
  string = paste('(', 1:len, ') ', string, sep = '')
  return(string)
}

data.0<-as_tibble(data.0)%>% 
  dplyr::mutate(temp = as.numeric(temperature))%>%
  dplyr::filter(rate>0)%>%
  dplyr::select(treatement,temp,rate)


nls_multstart_progress <- function(formula, data = parent.frame(), iter, start_lower, 
                                   start_upper, supp_errors = c("Y", "N"), convergence_count = 100, 
                                   control, modelweights, ...){
  if(!is.null(pb)){
    pb$tick()
  }
  nls_multstart(formula = formula, data = data, iter = iter, start_lower = start_lower, 
                start_upper = start_upper, supp_errors = supp_errors, convergence_count = convergence_count, 
                control = control, modelweights = modelweights, ...)
}

# start progress bar and estimate time it will take
number_of_models <- 1
number_of_curves <- length(unique(data.0$treatement))

# setup progress bar
pb <- progress::progress_bar$new(total = number_of_curves*number_of_models,
                                 clear = FALSE,
                                 format ="[:bar] :percent :elapsedfull")


d_fits <- nest(data.0, data = c(temp, rate)) %>%
  mutate(boatman = map(data, ~nls_multstart(rate~boatman_2017(temp = temp, rmax, tmin, tmax, a,b),
                                            data = .x,
                                            iter = c(4,4,4,4,4),
                                            start_lower = get_start_vals(.x$temp, .x$rate, model_name = 'boatman_2017') - 10,
                                            start_upper = get_start_vals(.x$temp, .x$rate, model_name = 'boatman_2017') + 10,
                                            lower = get_lower_lims(.x$temp, .x$rate, model_name = 'boatman_2017'),
                                            upper = get_upper_lims(.x$temp, .x$rate, model_name = 'boatman_2017'),
                                            supp_errors = 'Y',
                                            convergence_count = FALSE)))

#d_preds <- mutate(d_fits, new_data = map(data, ~tibble(temp = seq(min(.x$temp), max(.x$temp), length.out = 100)))) %>%
d_preds <- mutate(d_fits, new_data = map(data, ~tibble(temp = seq(16, 33, length.out = 100)))) %>%
  # get rid of original data column
  # select(., -data) %>%
  # stack models into a single column, with an id column for model_name
  pivot_longer(., names_to = 'model_name', values_to = 'fit', c(boatman))%>%
  # this uses both fit and new_data list columns
  mutate(preds = map2(fit, new_data, ~augment(.x, newdata = .y))) %>%
  # select only the columns we want to keep
  # unlist the preds list column
  unnest(preds)


# get coefs
d_fits <- mutate(d_fits, coefs = map(boatman, coef))


d_fits <- mutate(d_fits, nls_fit = map2(data, coefs, ~nlsLM(rate~boatman_2017(temp = temp, rmax, tmin, tmax, a,b),
                                                            data = .x,
                                                            start = .y,
                                                            lower = get_lower_lims(.x$temp, .x$rate, model_name = 'boatman_2017'),
                                                            upper = get_upper_lims(.x$temp, .x$rate, model_name = 'boatman_2017'))))


# create empty list column
d_fits <- mutate(d_fits, bootstrap = list(rep(NA, n())))

# run for loop to bootstrap each refitted model
for(i in 1:nrow(d_fits)){
  temp_data <- d_fits$data[[i]]
  temp_fit <- nlsLM(rate ~ boatman_2017(temp = temp, rmax, tmin, tmax, a,b),
                    data = temp_data,
                    start = d_fits$coefs[[i]])
  boot <- Boot(temp_fit, method = 'residual', R=1000)
  d_fits$bootstrap[[i]] <- boot
  rm(list = c('temp_fit', 'temp_data', 'boot'))
}


# get the raw values of each bootstrap
d_fits <- mutate(d_fits, output_boot = map(bootstrap, function(x) x$t))

# calculate predictions with a gnarly written function

d_fits <- mutate(d_fits, preds = map2(output_boot, data, function(x, y){
  temp <- as.data.frame(x) %>%
    drop_na() %>%
    mutate(iter = 1:n()) %>%
    group_by_all() %>%
    do(data.frame(temp = seq(min(y$temp), max(y$temp), length.out = 100))) %>%
    ungroup() %>%
    mutate(pred = boatman_2017(temp = temp, rmax, tmin, tmax, a,b))
  return(temp)
}))


data1<- read.table("fluct.txt", sep="\t", dec=",", header=TRUE)

data1<-as_tibble(data1)%>% 
  dplyr::mutate(temp=as.numeric(temp))%>%
  dplyr::mutate(g=as.numeric(g))%>%
  dplyr::mutate(lower=as.numeric(lower))%>%
  dplyr::mutate(upper=as.numeric(upper))%>%
  dplyr::select(treatement,type,g,lower,upper,temp)
data1$low<-data1$g-data1$lower

library(readxl)

data4<- read_excel("C:....../Supplementary_data/data/growth_rate_fluctuation.xlsx")
data4<-as_tibble(data4)%>% 
  dplyr::mutate(g = as.numeric(g))%>%
  dplyr::filter(Time==150)%>%
  dplyr::select(Treatment,Type,g)


data.0$Treatment<-'CT'
data.0<-as_tibble(data.0)%>% 
  dplyr::mutate(g = as.numeric(rate))%>%
  dplyr::mutate(Type = treatement)%>%
  dplyr::filter(temp==23.80)%>%
  dplyr::select(Treatment,Type,g)

data <- rbind(data4,data.0) 

data=data %>% mutate(treat=str_c(data$Treatment,data$Type,sep="-"))
# cr?ation de la fonction moyenne pour la fonction boot
moyenne<-function(data,indice)
{
  data.star <- data[indice]
  moy <- mean(data.star,na.rm=TRUE)
}

#permet de fixer la graine des tirages al?atoires et donc d'obtenir des r?sultats toujours identiques
set.seed(1234)

# realisation des ?chantillons bootstrap et estimation des moyennes
v.treat<- levels(as.factor(data$treat))
v.1.0 <- NULL
v.2.0 <- NULL
v.3.0 <- NULL

for (i in (1:length(v.treat))) {
  data2<- subset(data, treat == v.treat[i])  
  b1=boot(data2$g, statistic= moyenne, R=1000)
  
  ci=boot.ci(b1,type="perc")
  v<-c(ci[["percent"]])
  v.1<-v[4]
  v.1.0<-c(v.1.0,v.1)
  v.2<-v[5]
  v.2.0<-c(v.2.0,v.2)
  v.3<-mean(data2$g)
  v.3.0<-c(v.3.0,v.3)
  
}
tab.prms <- data.frame(v.treat, v.1.0,v.2.0,v.3.0)
tab.prms ##############g et indice 95
colnames(tab.prms)[colnames(tab.prms) == 'v.treat'] <- 'treat'
data.1=data1 %>% mutate(treat=str_c(data1$type,data1$treatement,sep="-"))
data.4<-left_join(data.1, tab.prms, by ='treat')

data.4<-as_tibble(data.4)%>% 
  dplyr::mutate(rate = as.numeric(v.3.0))%>%
  dplyr::mutate(lower = as.numeric(v.1.0))%>%
  dplyr::mutate(upper = as.numeric(v.2.0))%>%
  dplyr::select(treatement,type,rate,temp,lower,upper)

colnames(data.4)[colnames(data.4) == 'rate'] <- 'g'


library(readxl)
k<-8.62e-05

data.F12<- read_excel("C:..../Supplementary_data/data/F12.xlsx")
data.F24<- read_excel("C:..../Supplementary_data/data/F24.xlsx")
data.F48<- read_excel("C:..../Supplementary_data/data/F48.xlsx")
#############Y##########

#sharpeschoolhigh_1981


rate <- function(x){
  rmax * (sin(pi * ((x - tmin)/(tmax - tmin))^a))^b
 
}

library(MESS)#####AUC


v.rate.0 <- NULL
v.treat.0 <- NULL
v.Temp.0 <- NULL


v.int6.0<- NULL
###########i=food, j=repbootsrap########
for (i in c(1:3)){
  v.2<-i
  for(j in c(1:1000)){
    
    rmax<-  d_fits$output_boot[[i]][j,1]
    tmin<-  d_fits$output_boot[[i]][j,2]
    tmax<-  d_fits$output_boot[[i]][j,3]
    a<-  d_fits$output_boot[[i]][j,4]
    b<-  d_fits$output_boot[[i]][j,5]
    
    
    
    
    v.rate<-  (rmax * (sin(pi * ((data.F12$Temp - tmin)/(tmax - tmin))^a))^b)
    
    
    v.1<-mean(v.rate)
    v.4<-mean(data.F12$Temp)  
    v.rate.0 <- c(v.rate.0, v.1 )
    v.Temp.0 <- c(v.Temp.0, v.4 )
    v.treat.0<-c(v.treat.0,v.2)
    v.6<-auc(x=seq(1,length(v.rate),1),y=v.rate)/length(v.rate)
    v.int6.0 <- c(v.int6.0,v.6)     
    
  }
}


tab.prms.rate <- data.frame(v.rate.0, v.Temp.0,v.treat.0,v.int6.0) 

tab.prms.rate$v.treat.0[tab.prms.rate$v.treat.0==1]<-'Y'
tab.prms.rate$v.treat.0[tab.prms.rate$v.treat.0==2]<-'C'
tab.prms.rate$v.treat.0[tab.prms.rate$v.treat.0==3]<-'S'
tab.prms.rate$type<-'F12'

tab.prms.rate<-as_tibble(tab.prms.rate)%>% 
  dplyr::mutate(temp=as.numeric(v.Temp.0))%>%
  dplyr::mutate(g=as.numeric(v.int6.0))%>%
  dplyr::mutate(treatement=v.treat.0)%>%
  dplyr::select(treatement,g,temp,type)##########mean temperature = 23.5317

v.rate.0 <- NULL
v.treat.0 <- NULL
v.Temp.0 <- NULL
###########i=food, j=repbootsrap########
for (i in c(1:3)){
  v.2<-i
  for(j in c(1:1000)){
    
    rmax<-  d_fits$output_boot[[i]][j,1]
    tmin<-  d_fits$output_boot[[i]][j,2]
    tmax<-  d_fits$output_boot[[i]][j,3]
    a<-  d_fits$output_boot[[i]][j,4]
    b<-  d_fits$output_boot[[i]][j,5]
    
    
    
    
    v.rate<-  (rmax * (sin(pi * ((23.5317 - tmin)/(tmax - tmin))^a))^b)
    
    
  
    
    
    v.1<-v.rate
    v.4<-23.5317
    v.rate.0 <- c(v.rate.0, v.1 )
    v.Temp.0 <- c(v.Temp.0, v.4 )
    v.treat.0<-c(v.treat.0,v.2)
    
  }
}


tab.prms.rate.ctt <- data.frame(v.rate.0, v.Temp.0,v.treat.0) 

tab.prms.rate.ctt$v.treat.0[tab.prms.rate.ctt$v.treat.0==1]<-'Y'
tab.prms.rate.ctt$v.treat.0[tab.prms.rate.ctt$v.treat.0==2]<-'C'
tab.prms.rate.ctt$v.treat.0[tab.prms.rate.ctt$v.treat.0==3]<-'S'
tab.prms.rate.ctt$type<-'F12'

tab.prms.rate.ctt<-as_tibble(tab.prms.rate.ctt)%>% 
  dplyr::mutate(temp=as.numeric(v.Temp.0))%>%
  dplyr::mutate(g=as.numeric(v.rate.0))%>%
  dplyr::mutate(treatement=v.treat.0)%>%
  dplyr::select(treatement,g,temp,type)##########mean temperature = 23.5317

#############period 24##########


v.rate.0 <- NULL
v.treat.0 <- NULL
v.Temp.0 <- NULL

v.int6.0<- NULL


for (i in c(1:3)){
  v.2<-i
  for(j in c(1:1000)){
    
    rmax<-  d_fits$output_boot[[i]][j,1]
    tmin<-  d_fits$output_boot[[i]][j,2]
    tmax<-  d_fits$output_boot[[i]][j,3]
    a<-  d_fits$output_boot[[i]][j,4]
    b<-  d_fits$output_boot[[i]][j,5]
    
    
    
    
    v.rate<-  (rmax * (sin(pi * ((data.F24$Temp - tmin)/(tmax - tmin))^a))^b)
    
    v.1<-mean(v.rate)
    v.4<-mean(data.F24$Temp)  
    v.rate.0 <- c(v.rate.0, v.1 )
    v.Temp.0 <- c(v.Temp.0, v.4 )
    v.treat.0<-c(v.treat.0,v.2)
    
    v.6<-auc(x=seq(1,length(v.rate),1),y=v.rate)/length(v.rate)
    v.int6.0 <- c(v.int6.0,v.6)       
    
  }
}


tab.prms.rate.24 <- data.frame(v.rate.0, v.Temp.0,v.treat.0,v.int6.0) 

tab.prms.rate.24$v.treat.0[tab.prms.rate.24$v.treat.0==1]<-'Y'
tab.prms.rate.24$v.treat.0[tab.prms.rate.24$v.treat.0==2]<-'C'
tab.prms.rate.24$v.treat.0[tab.prms.rate.24$v.treat.0==3]<-'S'
tab.prms.rate.24$type<-'F24'

tab.prms.rate.24<-as_tibble(tab.prms.rate.24)%>% 
  dplyr::mutate(temp=as.numeric(v.Temp.0))%>%
  dplyr::mutate(g=as.numeric(v.int6.0))%>%
  dplyr::mutate(treatement=v.treat.0)%>%
  dplyr::select(treatement,g,temp,type)###########mean temperature =22.66641
v.rate.0 <- NULL
v.treat.0 <- NULL
v.Temp.0 <- NULL
###########i=food, j=repbootsrap########
for (i in c(1:3)){
  v.2<-i
  for(j in c(1:1000)){
    
    rmax<-  d_fits$output_boot[[i]][j,1]
    tmin<-  d_fits$output_boot[[i]][j,2]
    tmax<-  d_fits$output_boot[[i]][j,3]
    a<-  d_fits$output_boot[[i]][j,4]
    b<-  d_fits$output_boot[[i]][j,5]
    
    
    
    
    v.rate<-  (rmax * (sin(pi * ((22.66641- tmin)/(tmax - tmin))^a))^b)
    
   
    
    v.1<-v.rate
    v.4<-22.66641
    v.rate.0 <- c(v.rate.0, v.1 )
    v.Temp.0 <- c(v.Temp.0, v.4 )
    v.treat.0<-c(v.treat.0,v.2)
    
  }
}


tab.prms.rate.24.ctt <- data.frame(v.rate.0, v.Temp.0,v.treat.0) 

tab.prms.rate.24.ctt$v.treat.0[tab.prms.rate.24.ctt$v.treat.0==1]<-'Y'
tab.prms.rate.24.ctt$v.treat.0[tab.prms.rate.24.ctt$v.treat.0==2]<-'C'
tab.prms.rate.24.ctt$v.treat.0[tab.prms.rate.24.ctt$v.treat.0==3]<-'S'
tab.prms.rate.24.ctt$type<-'F24'

tab.prms.rate.24.ctt<-as_tibble(tab.prms.rate.24.ctt)%>% 
  dplyr::mutate(temp=as.numeric(v.Temp.0))%>%
  dplyr::mutate(g=as.numeric(v.rate.0))%>%
  dplyr::mutate(treatement=v.treat.0)%>%
  dplyr::select(treatement,g,temp,type)

#####################period48###########

v.rate.0 <- NULL
v.treat.0 <- NULL
v.Temp.0 <- NULL


v.int6.0<- NULL

for (i in c(1:3)){
  v.2<-i
  for(j in c(1:1000)){
    
    rmax<-  d_fits$output_boot[[i]][j,1]
    tmin<-  d_fits$output_boot[[i]][j,2]
    tmax<-  d_fits$output_boot[[i]][j,3]
    a<-  d_fits$output_boot[[i]][j,4]
    b<-  d_fits$output_boot[[i]][j,5]
    
    
    
    
    v.rate<-  (rmax * (sin(pi * ((data.F48$Temp - tmin)/(tmax - tmin))^a))^b)
    v.1<-mean(v.rate)
    v.4<-mean(data.F48$Temp)  
    v.rate.0 <- c(v.rate.0, v.1 )
    v.Temp.0 <- c(v.Temp.0, v.4 )
    v.treat.0<-c(v.treat.0,v.2)
    v.6<-auc(x=seq(1,length(v.rate),1),y=v.rate)/length(v.rate)
    v.int6.0 <- c(v.int6.0,v.6) 
  }
}


tab.prms.rate.48 <- data.frame(v.rate.0, v.Temp.0,v.treat.0,v.int6.0) 

tab.prms.rate.48$v.treat.0[tab.prms.rate.48$v.treat.0==1]<-'Y'
tab.prms.rate.48$v.treat.0[tab.prms.rate.48$v.treat.0==2]<-'C'
tab.prms.rate.48$v.treat.0[tab.prms.rate.48$v.treat.0==3]<-'S'
tab.prms.rate.48$type<-'F48'


tab.prms.rate.48<-as_tibble(tab.prms.rate.48)%>% 
  dplyr::mutate(temp=as.numeric(v.Temp.0))%>%
  dplyr::mutate(g=as.numeric(v.int6.0))%>%
  dplyr::mutate(treatement=v.treat.0)%>%
  dplyr::select(treatement,g,temp,type)####mean temperature =22.68369
v.rate.0 <- NULL
v.treat.0 <- NULL
v.Temp.0 <- NULL
###########i=food, j=repbootsrap########
for (i in c(1:3)){
  v.2<-i
  for(j in c(1:1000)){
    
    rmax<-  d_fits$output_boot[[i]][j,1]
    tmin<-  d_fits$output_boot[[i]][j,2]
    tmax<-  d_fits$output_boot[[i]][j,3]
    a<-  d_fits$output_boot[[i]][j,4]
    b<-  d_fits$output_boot[[i]][j,5]
    
    
    
    
    v.rate<-  (rmax * (sin(pi * ((22.68369 - tmin)/(tmax - tmin))^a))^b)
    
   
    
    
    v.1<-v.rate
    v.4<-22.68369
    v.rate.0 <- c(v.rate.0, v.1 )
    v.Temp.0 <- c(v.Temp.0, v.4 )
    v.treat.0<-c(v.treat.0,v.2)
    
  }
}


tab.prms.rate.48.ctt <- data.frame(v.rate.0, v.Temp.0,v.treat.0) 

tab.prms.rate.48.ctt$v.treat.0[tab.prms.rate.48.ctt$v.treat.0==1]<-'Y'
tab.prms.rate.48.ctt$v.treat.0[tab.prms.rate.48.ctt$v.treat.0==2]<-'C'
tab.prms.rate.48.ctt$v.treat.0[tab.prms.rate.48.ctt$v.treat.0==3]<-'S'
tab.prms.rate.48.ctt$type<-'F48'

tab.prms.rate.48.ctt<-as_tibble(tab.prms.rate.48.ctt)%>% 
  dplyr::mutate(temp=as.numeric(v.Temp.0))%>%
  dplyr::mutate(g=as.numeric(v.rate.0))%>%
  dplyr::mutate(treatement=v.treat.0)%>%
  dplyr::select(treatement,g,temp,type)


data_freq<-rbind(tab.prms.rate,tab.prms.rate.24,tab.prms.rate.48)
data_freq_ctt<-rbind(tab.prms.rate.ctt,tab.prms.rate.24.ctt,tab.prms.rate.48.ctt)

pd=position_dodge(0.3)

dataf<- read_excel("C:....../Supplementary_data/data/growth_rate_fluctuation.xlsx")

dataf<-dataf[-which(dataf$Treatment =='cst'),]
dataf= dataf %>% filter(Time=="150")
dataf<-as_tibble(dataf)%>% 
  dplyr::mutate(g=as.numeric(g))%>%
  dplyr::mutate(type=Treatment)%>%
  dplyr::mutate(treatement=Type)%>%
  dplyr::select(treatement,g,type)

data_freq<-as_tibble(data_freq)%>% 
  dplyr::select(treatement,g,type)
data_freq_ctt<-as_tibble(data_freq_ctt)%>% 
  dplyr::select(treatement,g,type)

dataf$class<- 'obs'
data_freq$class<-'pred'
data_freq_ctt$class<-'ctt'
data<-rbind(dataf,data_freq,data_freq_ctt)

data=data %>% mutate(treat=str_c(data$treatement,data$type,data$class,sep="-"))
# cr?ation de la fonction moyenne pour la fonction boot
moyenne<-function(data,indice)
{
  data.star <- data[indice]
  moy <- mean(data.star,na.rm=TRUE)
}

#permet de fixer la graine des tirages al?atoires et donc d'obtenir des r?sultats toujours identiques
set.seed(1234)

# realisation des ?chantillons bootstrap et estimation des moyennes
v.treat<- levels(as.factor(data$treat))
v.1.0 <- NULL
v.2.0 <- NULL
v.3.0 <- NULL
v.4.0 <- NULL
v.5.0 <- NULL
v.6.0 <- NULL
for (i in (1:length(v.treat))) {
  data2<- subset(data, treat == v.treat[i])  
  b1=boot(data2$g, statistic= moyenne, R=1000)
  
  ci=boot.ci(b1,type="perc")
  v<-c(ci[["percent"]])
  v.1<-v[4]
  v.1.0<-c(v.1.0,v.1)
  v.2<-v[5]
  v.2.0<-c(v.2.0,v.2)
  v.3<-mean(data2$g)
  v.3.0<-c(v.3.0,v.3)
  v.4<-first(data2$treatement)
  v.4.0<-c(v.4.0,v.4)
  v.5<-first(data2$type)
  v.5.0<-c(v.5.0,v.5)
  v.6<-first(data2$class)
  v.6.0<-c(v.6.0,v.6)
}
tab.prms <- data.frame(v.treat, v.1.0,v.2.0,v.3.0,v.4.0,v.5.0,v.6.0)
tab.prms ##############g et indice 95
colnames(tab.prms)[colnames(tab.prms) == 'v.treat'] <- 'treat'


data<-as_tibble(tab.prms)%>% 
  dplyr::mutate(rate = as.numeric(v.3.0))%>%
  dplyr::mutate(lower = as.numeric(v.1.0))%>%
  dplyr::mutate(upper = as.numeric(v.2.0))%>%
  dplyr::mutate(treatement = v.4.0)%>%
  dplyr::mutate(type = v.5.0)%>%
  dplyr::mutate(class = v.6.0)%>%
  dplyr::select(treatement,type,rate,class,lower,upper)

colnames(data)[colnames(data) == 'rate'] <- 'g'


pd=position_dodge(0) 


data.Y<-as_tibble(data)%>% 
  dplyr::filter(treatement=="Y")
data.Y$type[data.Y$type=='F12']<-12
data.Y$type[data.Y$type=='F24']<-24
data.Y$type[data.Y$type=='F48']<-48
a<-ggplot(data.Y,aes(x=type, y=g,group = class,col=class),show.legend = FALSE) +
  geom_errorbar(aes(x=type,y=g,ymin=lower,ymax=upper),data.Y,fill=class, width=.1, size=0.15,show.legend = FALSE,  position=pd)+
  geom_point(size=3,show.legend = FALSE)+
  geom_line(size=1,show.legend = FALSE)+
  geom_point(shape = 1,size = 3,colour = "black")+
  theme(  panel.background = element_blank(),
          panel.grid.major = element_blank(), 
          panel.grid.minor = element_blank(),
          panel.border = element_rect(colour = "black",fill=NA))+
  scale_color_manual(breaks = c("obs", "pred","ctt"),values=c("cyan3","black","#ffec1b"),
                     labels=c("Observation","Prediction","Constant"))+
  
  labs(x = '',
       y = 'Growth rate (d-1)',title='')+
  theme(axis.text.x = element_text(size=8),axis.text.y = element_text(size=8),
        axis.title.x = element_text(size=10),axis.title.y = element_text(size=10))+ylim(0.40,0.65)


data.C<-as_tibble(data)%>% 
  dplyr::filter(treatement=="C")
data.C$type[data.C$type=='F12']<-12
data.C$type[data.C$type=='F24']<-24
data.C$type[data.C$type=='F48']<-48
b<-ggplot(data.C,aes(x=type, y=g,group = class,col=class),show.legend = FALSE) + 
  geom_errorbar(aes(x=type,y=g,ymin=lower,ymax=upper),data.C,fill=class, width=.1, size=0.15,show.legend = FALSE,  position=pd)+
  geom_point(size=3,show.legend = FALSE)+
  geom_line(size=1,show.legend = FALSE)+
  geom_point(shape = 1,size = 3,colour = "black")+
  theme(  panel.background = element_blank(),
          panel.grid.major = element_blank(), 
          panel.grid.minor = element_blank(),
          panel.border = element_rect(colour = "black",fill=NA))+
  scale_color_manual(breaks = c("obs", "pred","ctt"),values=c("cyan3","black","#ffec1b"),
                     labels=c("Observation","Prediction","Constant"))+
  
  labs(x = 'Thermal Fluctuation period (h)',
       y = '',title='')+
  theme(axis.text.x = element_text(size=8),axis.text.y = element_text(size=8),
        axis.title.x = element_text(size=10),axis.title.y = element_text(size=10))+ylim(0.40,0.65)

data.S<-as_tibble(data)%>% 
  dplyr::filter(treatement=="S")
data.S$type[data.S$type=='F12']<-12
data.S$type[data.S$type=='F24']<-24
data.S$type[data.S$type=='F48']<-48
c<-ggplot(data.S,aes(x=type, y=g,group = class,col=class),show.legend = FALSE) +
  geom_errorbar(aes(x=type,y=g,ymin=lower,ymax=upper),data.S,fill=class, width=.1, size=0.15,show.legend = FALSE,  position=pd)+
  geom_point(size=3,show.legend = FALSE)+
  geom_line(size=1,show.legend = FALSE)+
  geom_point(shape = 1,size = 3,colour = "black")+
  theme(  panel.background = element_blank(),
          panel.grid.major = element_blank(), 
          panel.grid.minor = element_blank(),
          panel.border = element_rect(colour = "black",fill=NA))+
  scale_color_manual(breaks = c("obs", "pred","ctt"),values=c("orchid","black","#ffec1b"),
                     labels=c("Observation","Prediction","Constant"))+
  labs(x = '',
       y = '',title='')+
  theme(axis.text.x = element_text(size=8),axis.text.y = element_text(size=8),
        axis.title.x = element_text(size=10),axis.title.y = element_text(size=10))+ylim(0.40,0.65)

library("cowplot")
plot_grid(a,b,c ,ncol = 3, nrow = 1)


####################################


data.mean=data %>% mutate(treat=str_c(data$type,data$class,data$treatement,sep="-"))
data.mean=data.mean %>% group_by(treat) %>% summarise(treatement=first(treatement),class= first(class),type=first(type),
                                                      g=mean(g))

library(MKinfer)
pred.C.fast.f<-data_freq %>% filter(type=="F12" & treatement== "C")
obs.C.fast.f <-dataf %>% filter(type=="F12" & treatement== "C")
ctt.C.fast.f<-data_freq_ctt %>% filter(type=="F12" & treatement== "C")

pred.C.mid.f<-data_freq %>% filter(type=="F24" & treatement== "C")
obs.C.mid.f <-dataf %>% filter(type=="F24" & treatement== "C")
ctt.C.mid.f<-data_freq_ctt %>% filter(type=="F24" & treatement== "C")

pred.C.slow.f<-data_freq %>% filter(type=="F48" & treatement== "C")
obs.C.slow.f <-dataf %>% filter(type=="F48" & treatement== "C")
ctt.C.slow.f<-data_freq_ctt %>% filter(type=="F48" & treatement== "C")

pred.Y.fast.f<-data_freq %>% filter(type=="F12" & treatement== "Y")
obs.Y.fast.f <-dataf %>% filter(type=="F12" & treatement== "Y")
ctt.Y.fast.f<-data_freq_ctt %>% filter(type=="F12" & treatement== "Y")

pred.Y.mid.f<-data_freq %>% filter(type=="F24" & treatement== "Y")
obs.Y.mid.f <-dataf %>% filter(type=="F24" & treatement== "Y")
ctt.Y.mid.f<-data_freq_ctt %>% filter(type=="F24" & treatement== "Y")

pred.Y.slow.f<-data_freq %>% filter(type=="F48" & treatement== "Y")
obs.Y.slow.f <-dataf %>% filter(type=="F48" & treatement== "Y")
ctt.Y.slow.f<-data_freq_ctt %>% filter(type=="F48" & treatement== "Y")

pred.S.fast.f<-data_freq %>% filter(type=="F12" & treatement== "S")
obs.S.fast.f <-dataf %>% filter(type=="F12" & treatement== "S")
ctt.S.fast.f<-data_freq_ctt %>% filter(type=="F12" & treatement== "S")

pred.S.mid.f<-data_freq %>% filter(type=="F24" & treatement== "S")
obs.S.mid.f <-dataf %>% filter(type=="F24" & treatement== "S")
ctt.S.mid.f<-data_freq_ctt %>% filter(type=="F24" & treatement== "S")

pred.S.slow.f<-data_freq %>% filter(type=="F48" & treatement== "S")
obs.S.slow.f <-dataf %>% filter(type=="F48" & treatement== "S")
ctt.S.slow.f<-data_freq_ctt %>% filter(type=="F48" & treatement== "S")

boot.t.test(x=pred.Y.slow.f$g, y=obs.Y.slow.f$g)
boot.t.test(x=pred.Y.mid.f$g, y=obs.Y.mid.f$g)
boot.t.test(x=pred.Y.fast.f$g, y=obs.Y.fast.f$g)

boot.t.test(x=pred.C.slow.f$g, y=obs.C.slow.f$g)
boot.t.test(x=pred.C.mid.f$g, y=obs.C.mid.f$g)
boot.t.test(x=pred.C.fast.f$g, y=obs.C.fast.f$g)

boot.t.test(x=pred.S.slow.f$g, y=obs.S.slow.f$g)
boot.t.test(x=pred.S.mid.f$g, y=obs.S.mid.f$g)
boot.t.test(x=pred.S.fast.f$g, y=obs.S.fast.f$g)


boot.t.test(x=ctt.Y.slow.f$g, y=obs.Y.slow.f$g)
boot.t.test(x=ctt.Y.mid.f$g, y=obs.Y.mid.f$g)
boot.t.test(x=ctt.Y.fast.f$g, y=obs.Y.fast.f$g)

boot.t.test(x=ctt.C.slow.f$g, y=obs.C.slow.f$g)
boot.t.test(x=ctt.C.mid.f$g, y=obs.C.mid.f$g)
boot.t.test(x=ctt.C.fast.f$g, y=obs.C.fast.f$g)

boot.t.test(x=ctt.S.slow.f$g, y=obs.S.slow.f$g)
boot.t.test(x=ctt.S.mid.f$g, y=obs.S.mid.f$g)
boot.t.test(x=ctt.S.fast.f$g, y=obs.S.fast.f$g)
```
