## Supplementary material for "Diet quality alters the consequences of gradual phenotypic plasticity for individual performance": Data and Script: model_comparaison.R

```
library(boot)
library(car)
library(rTPC)
library(nls.multstart)
library(broom)
library(tidyverse)
library(patchwork)
library(minpack.lm)
library(tis)
library(pROC)
setwd("C:.../Supplementary_data/data")
data.0<- read.table("growth_rate_constant.txt", sep="\t", dec=",", header=TRUE)

data.0$M<-data.0$M*1000####mass in mg
mi=8.8##############mean initial mass
data.0$rate<-(log(data.0$M)-log(mi))/(data.0$time*0.0416667)


knitr::opts_chunk$set(
  collapse = TRUE,
  comment = "#>",
  #tidy.opts=list(width.cutoff=60),
  #tidy=TRUE,
  fig.align = 'center'
)
data.0<-as_tibble(data.0)%>% 
  dplyr::mutate(temp = as.numeric(temperature))%>%
  dplyr::mutate(day=as.numeric(day))%>%
  dplyr::mutate(hour=as.numeric(time))%>%
  dplyr::filter(rate>0)%>%
  dplyr::select(treatement,temp,rate)

###############
data.Y<-as_tibble(data.0)%>% 
  dplyr::filter(treatement=="Y")%>%
  dplyr::select(temp,rate)

data.C<-as_tibble(data.0)%>% 
  dplyr::filter(treatement=="C")%>%
  dplyr::select(temp,rate)

data.S<-as_tibble(data.0)%>% 
  dplyr::filter(treatement=="S")%>%
  dplyr::select(temp,rate)

d_fits_S <- nest(data.S, data = c(temp, rate)) %>%
            mutate(boatman = map(data, ~nls_multstart(rate~boatman_2017(temp = temp, rmax, tmin, tmax, a,b),
                                                      data = .x,
                                                      iter = c(4,4,4,4,4),
                                                      start_lower = get_start_vals(.x$temp, .x$rate, model_name = 'boatman_2017') - 10,
                                                      start_upper = get_start_vals(.x$temp, .x$rate, model_name = 'boatman_2017') + 10,
                                                      lower = get_lower_lims(.x$temp, .x$rate, model_name = 'boatman_2017'),
                                                      upper = get_upper_lims(.x$temp, .x$rate, model_name = 'boatman_2017'),
                                                      supp_errors = 'Y',
                                                      convergence_count = FALSE)),
                   pawar = map(data, ~nls_multstart(rate~pawar_2018(temp = temp, r_tref, e, eh, topt, tref = 15),
                                                    data = .x,
                                                    iter = c(4,4,4,4),
                                                    start_lower = get_start_vals(.x$temp, .x$rate, model_name = 'pawar_2018') - 10,
                                                    start_upper = get_start_vals(.x$temp, .x$rate, model_name = 'pawar_2018') + 10,
                                                    lower = get_lower_lims(.x$temp, .x$rate, model_name = 'pawar_2018'),
                                                    upper = get_upper_lims(.x$temp, .x$rate, model_name = 'pawar_2018'),
                                                    supp_errors = 'Y',
                                                    convergence_count = FALSE)),
                   beta = map(data, ~nls_multstart(rate~beta_2012(temp = temp, a, b, c, d, e),
                                                   data = .x,
                                                   iter = c(6,6,6,6,6),
                                                   start_lower = get_start_vals(.x$temp, .x$rate, model_name = 'beta_2012') - 10,
                                                   start_upper = get_start_vals(.x$temp, .x$rate, model_name = 'beta_2012') + 10,
                                                   lower = get_lower_lims(.x$temp, .x$rate, model_name = 'beta_2012'),
                                                   upper = get_upper_lims(.x$temp, .x$rate, model_name = 'beta_2012'),
                                                   supp_errors = 'Y',
                                                   convergence_count = FALSE)),
                   briere2 = map(data, ~nls_multstart(rate~briere2_1999(temp = temp, tmin, tmax, a,b),
                                                      data = .x,
                                                      iter = c(4,4,4,4),
                                                      start_lower = get_start_vals(.x$temp, .x$rate, model_name = 'briere2_1999') - 10,
                                                      start_upper = get_start_vals(.x$temp, .x$rate, model_name = 'briere2_1999') + 10,
                                                      lower = get_lower_lims(.x$temp, .x$rate, model_name = 'briere2_1999'),
                                                      upper = get_upper_lims(.x$temp, .x$rate, model_name = 'briere2_1999'),
                                                      supp_errors = 'Y',
                                                      convergence_count = FALSE)),
                   gaussian = map(data, ~nls_multstart(rate~gaussian_1987(temp = temp, rmax, topt, a),
                                                       data = .x,
                                                       iter = c(4,4,4),
                                                       start_lower = get_start_vals(.x$temp, .x$rate, model_name = 'gaussian_1987') - 10,
                                                       start_upper = get_start_vals(.x$temp, .x$rate, model_name = 'gaussian_1987') + 10,
                                                       lower = get_lower_lims(.x$temp, .x$rate, model_name = 'gaussian_1987'),
                                                       upper = get_upper_lims(.x$temp, .x$rate, model_name = 'gaussian_1987'),
                                                       supp_errors = 'Y',
                                                       convergence_count = FALSE)),
                   quadratic = map(data, ~nls_multstart(rate~quadratic_2008(temp = temp, a, b, c),
                                                        data = .x,
                                                        iter = c(4,4,4),
                                                        start_lower = get_start_vals(.x$temp, .x$rate, model_name = 'quadratic_2008') - 0.5,
                                                        start_upper = get_start_vals(.x$temp, .x$rate, model_name = 'quadratic_2008') + 0.5,
                                                        lower = get_lower_lims(.x$temp, .x$rate, model_name = 'quadratic_2008'),
                                                        upper = get_upper_lims(.x$temp, .x$rate, model_name = 'quadratic_2008'),
                                                        supp_errors = 'Y',
                                                        convergence_count = FALSE)),
                   weibull = map(data, ~nls_multstart(rate~weibull_1995(temp = temp, a,topt,b,c),
                                                      data = .x,
                                                      iter = c(4,4,4,4),
                                                      start_lower = get_start_vals(.x$temp, .x$rate, model_name = 'weibull_1995') - 10,
                                                      start_upper = get_start_vals(.x$temp, .x$rate, model_name = 'weibull_1995') + 10,
                                                      lower = get_lower_lims(.x$temp, .x$rate, model_name = 'weibull_1995'),
                                                      upper = get_upper_lims(.x$temp, .x$rate, model_name = 'weibull_1995'),
                                                      supp_errors = 'Y',
                                                      convergence_count = FALSE)),
                   thomas1 = map(data, ~nls_multstart(rate~thomas_2012(temp = temp, a,b,c,topt),
                                                      data = .x,
                                                      iter = c(4,4,4,4),
                                                      start_lower = get_start_vals(.x$temp, .x$rate, model_name = 'thomas_2012') - 1,
                                                      start_upper = get_start_vals(.x$temp, .x$rate, model_name = 'thomas_2012') + 2,
                                                      lower = get_lower_lims(.x$temp, .x$rate, model_name = 'thomas_2012'),
                                                      upper = get_upper_lims(.x$temp, .x$rate, model_name = 'thomas_2012'),
                                                      supp_errors = 'Y',
                                                      convergence_count = FALSE)),
                   lactin2 = map(data, ~nls_multstart(rate~lactin2_1995(temp = temp, a, b, tmax, delta_t),
                                                      data = .x,
                                                      iter = c(4,4,4,4),
                                                      start_lower = get_start_vals(.x$temp, .x$rate, model_name = 'lactin2_1995') - 10,
                                                      start_upper = get_start_vals(.x$temp, .x$rate, model_name = 'lactin2_1995') + 10,
                                                      lower = get_lower_lims(.x$temp, .x$rate, model_name = 'lactin2_1995'),
                                                      upper = get_upper_lims(.x$temp, .x$rate, model_name = 'lactin2_1995'),
                                                      supp_errors = 'Y',
                                                      convergence_count = FALSE)),
          sharpeschoolhigh = map(data, ~nls_multstart(rate~sharpeschoolhigh_1981(temp = temp, r_tref,e,eh,th, tref = 15),
                                                     data = .x,
                                                     iter = c(4,4,4,4),
                                                     start_lower = get_start_vals(.x$temp, .x$rate, model_name = 'sharpeschoolhigh_1981') - 10,
                                                     start_upper = get_start_vals(.x$temp, .x$rate, model_name = 'sharpeschoolhigh_1981') + 10,
                                                     lower = get_lower_lims(.x$temp, .x$rate, model_name = 'sharpeschoolhigh_1981'),
                                                     upper = get_upper_lims(.x$temp, .x$rate, model_name = 'sharpeschoolhigh_1981'),
                                                     supp_errors = 'Y',
                                                     convergence_count = FALSE)))


################
# stack models
d_stack_Y <- select(d_fits_Y, -data) %>%
  pivot_longer(., names_to = 'model_name', values_to = 'fit', boatman:sharpeschoolhigh)

# get predictions using augment
newdata<- tibble(temp = seq(min(data.Y$temp), max(data.Y$temp), length.out = 100))
d_preds_Y <- d_stack_Y %>%
  mutate(., preds = map(fit, augment, newdata = newdata)) %>%
  select(-fit) %>%
  unnest(preds)

# take a random point from each model for labelling
d_labs_Y <- filter(d_preds_Y, temp < 30) %>%
  group_by(., model_name) %>%
  sample_n(., 1) %>%
  ungroup()
library(ggrepel)
# plot
ggplot(d_preds_Y, aes(temp, .fitted)) +
  geom_line(aes(col = model_name)) +
  geom_label_repel(aes(temp, .fitted, label = model_name, col = model_name), fill = 'white', nudge_y = 0.8, segment.size = 0.2, segment.colour = 'grey50', d_labs_Y) +
  geom_point(aes(temp, rate), data.Y) +
  theme_bw(base_size = 12) +
  theme(legend.position = 'none') +
  labs(x = 'Temperature (?C)',
       y = 'rate',
       title = '') +
  geom_hline(aes(yintercept = 0), linetype = 2) +
  scale_color_brewer(type = 'qual', palette = 2)


d_ic_Y <- d_stack_Y %>%
  mutate(., info = map(fit, glance),
         AICc =  map_dbl(fit, MuMIn::AICc)) %>%
  select(-fit) %>%
  unnest(info) %>%
  select(model_name, sigma, AIC, AICc, BIC, df.residual)

d_ic_Y


best_model_Y = filter(d_ic_Y, AICc == min(AICc)) %>% pull(model_name)
best_model_Y
col_best_mod_Y = RColorBrewer::brewer.pal(n = 6, name = "Dark2")[6]

# plot
ggplot(d_preds_Y, aes(temp, .fitted)) +
  geom_line(aes(group = model_name), col = 'grey50', alpha = 0.5) +
  geom_line(data = filter(d_preds_Y, model_name == best_model_Y), col = col_best_mod_Y) +
  geom_label_repel(aes(temp, .fitted, label = model_name), fill = 'white', nudge_y = 0.8, segment.size = 0.2, segment.colour = 'grey50', data = filter(d_labs_Y, model_name == best_model_Y), col = col_best_mod_Y) +
  geom_point(aes(temp, rate), data.Y) +
  theme_bw(base_size = 12) +
  theme(legend.position = 'none') +
  labs(x = 'Temperature (?C)',
       y = ' rate',
       title = '',
       subtitle= 'best model') +
  geom_hline(aes(yintercept = 0), linetype = 2) 


# stack models
d_stack_C <- select(d_fits_C, -data) %>%
  pivot_longer(., names_to = 'model_name', values_to = 'fit', boatman:sharpeschoolhigh)

# get predictions using augment
newdata<- tibble(temp = seq(min(data.C$temp), max(data.C$temp), length.out = 100))
d_preds_C <- d_stack_C %>%
  mutate(., preds = map(fit, augment, newdata = newdata)) %>%
  select(-fit) %>%
  unnest(preds)

# take a random point from each model for labelling
d_labs_C <- filter(d_preds_C, temp < 30) %>%
  group_by(., model_name) %>%
  sample_n(., 1) %>%
  ungroup()
library(ggrepel)
# plot
ggplot(d_preds_C, aes(temp, .fitted)) +
  geom_line(aes(col = model_name)) +
  geom_label_repel(aes(temp, .fitted, label = model_name, col = model_name), fill = 'white', nudge_y = 0.8, segment.size = 0.2, segment.colour = 'grey50', d_labs_C) +
  geom_point(aes(temp, rate), data.C) +
  theme_bw(base_size = 12) +
  theme(legend.position = 'none') +
  labs(x = 'Temperature (?C)',
       y = 'rate',
       title = '') +
  geom_hline(aes(yintercept = 0), linetype = 2) +
  scale_color_brewer(type = 'qual', palette = 2)


d_ic_C <- d_stack_C %>%
  mutate(., info = map(fit, glance),
         AICc =  map_dbl(fit, MuMIn::AICc)) %>%
  select(-fit) %>%
  unnest(info) %>%
  select(model_name, sigma, AIC, AICc, BIC, df.residual)

d_ic_C


best_model_C = filter(d_ic_C, AICc == min(AICc)) %>% pull(model_name)
best_model_C
col_best_mod_C = RColorBrewer::brewer.pal(n = 6, name = "Dark2")[6]

# plot
ggplot(d_preds_C, aes(temp, .fitted)) +
  geom_line(aes(group = model_name), col = 'grey50', alpha = 0.5) +
  geom_line(data = filter(d_preds_C, model_name == best_model_C), col = col_best_mod_C) +
  geom_label_repel(aes(temp, .fitted, label = model_name), fill = 'white', nudge_y = 0.8, segment.size = 0.2, segment.colour = 'grey50', data = filter(d_labs_C, model_name == best_model_C), col = col_best_mod_C) +
  geom_point(aes(temp, rate), data.C) +
  theme_bw(base_size = 12) +
  theme(legend.position = 'none') +
  labs(x = 'Temperature (?C)',
       y = ' rate',
       title = '',
       subtitle= 'best model') +
  geom_hline(aes(yintercept = 0), linetype = 2) 


# stack models
d_stack_S <- select(d_fits_S, -data) %>%
  pivot_longer(., names_to = 'model_name', values_to = 'fit', boatman:sharpeschoolhigh)

# get predictions using augment
newdata<- tibble(temp = seq(min(data.S$temp), max(data.S$temp), length.out = 100))
d_preds_S <- d_stack_S %>%
  mutate(., preds = map(fit, augment, newdata = newdata)) %>%
  select(-fit) %>%
  unnest(preds)

# take a random point from each model for labelling
d_labs_S <- filter(d_preds_S, temp < 30) %>%
  group_by(., model_name) %>%
  sample_n(., 1) %>%
  ungroup()
library(ggrepel)
# plot
ggplot(d_preds_S, aes(temp, .fitted)) +
  geom_line(aes(col = model_name)) +
  geom_label_repel(aes(temp, .fitted, label = model_name, col = model_name), fill = 'white', nudge_y = 0.8, segment.size = 0.2, segment.colour = 'grey50', d_labs_S) +
  geom_point(aes(temp, rate), data.S) +
  theme_bw(base_size = 12) +
  theme(legend.position = 'none') +
  labs(x = 'Temperature (?C)',
       y = 'rate',
       title = '') +
  geom_hline(aes(yintercept = 0), linetype = 2) +
  scale_color_brewer(type = 'qual', palette = 2)


d_ic_S <- d_stack_S %>%
  mutate(., info = map(fit, glance),
         AICc =  map_dbl(fit, MuMIn::AICc)) %>%
  select(-fit) %>%
  unnest(info) %>%
  select(model_name, sigma, AIC, AICc, BIC, df.residual)

d_ic_S


best_model_S = filter(d_ic_S, AICc == min(AICc)) %>% pull(model_name)
best_model_S
col_best_mod_S = RColorBrewer::brewer.pal(n = 6, name = "Dark2")[6]

# plot
ggplot(d_preds_S, aes(temp, .fitted)) +
  geom_line(aes(group = model_name), col = 'grey50', alpha = 0.5) +
  geom_line(data = filter(d_preds_S, model_name == best_model_S), col = col_best_mod_S) +
  geom_label_repel(aes(temp, .fitted, label = model_name), fill = 'white', nudge_y = 0.8, segment.size = 0.2, segment.colour = 'grey50', data = filter(d_labs_C, model_name == best_model_S), col = col_best_mod_S) +
  geom_point(aes(temp, rate), data.S) +
  theme_bw(base_size = 12) +
  theme(legend.position = 'none') +
  labs(x = 'Temperature (?C)',
       y = ' rate',
       title = '',
       subtitle= ' best model') +
  geom_hline(aes(yintercept = 0), linetype = 2) 

d_ic_S
d_ic_Y
d_ic_C
##############
```
