## Supplementary material for "Diet quality alters the consequences of gradual phenotypic plasticity for individual performance": Data and Script: Models.pdf

**Figure S1: Concurrent thermal performance curve (TPC) model fits** for *Daphnia magna* growth rate ($d^{-1}$) under various constant temperatures. The most parsimonious models based on the AICc score are shown in yellow.

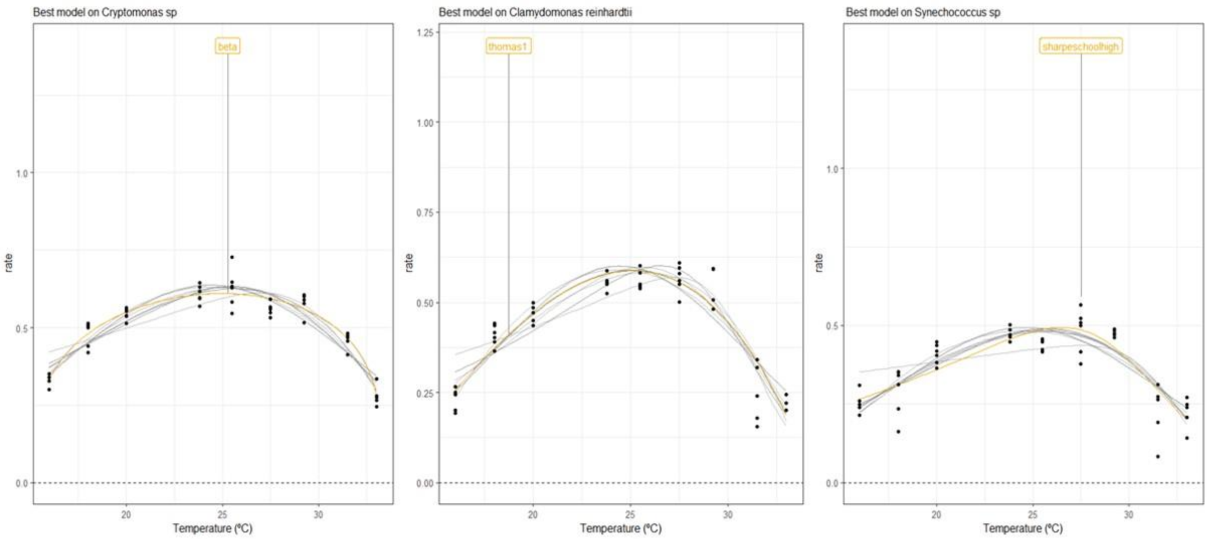

**Figure S2:** Daphnia magna growth rate ($day^{-1}$) under different diets ((A) *Cryptomonas sp*., (B) *Chlamydomonas reinhardtii*., (C) *Synechococcus sp*.) for each thermal period treatment using the different concurrent TPC models (see Figure S1) ((A) Briere2, (B) Boatman 2017, (C) Beta 2012, (D) Gaussian 1987, (E) Lactin 1995, (F) Quadratic 2008, (G) Sharpeschoolhigh 1981, (H) Thomas 2012, (I) Weibull 1995 and (J) Pawar 2018). The equations of the models can be found in supplementary data (models) (Fey et al. 2021) **Observed growth rate** $\overline{P(T,\Phi)}$ (*blue*: growth rate lower than expected, *pink*: growth rate higher than expected), Theoretical **time-averaged performance** $\overline{P(T,\Phi^*)}$ (*black*) and the **mean temperature performance** $P(\bar{T},\Phi^*)$ (*yellow*).

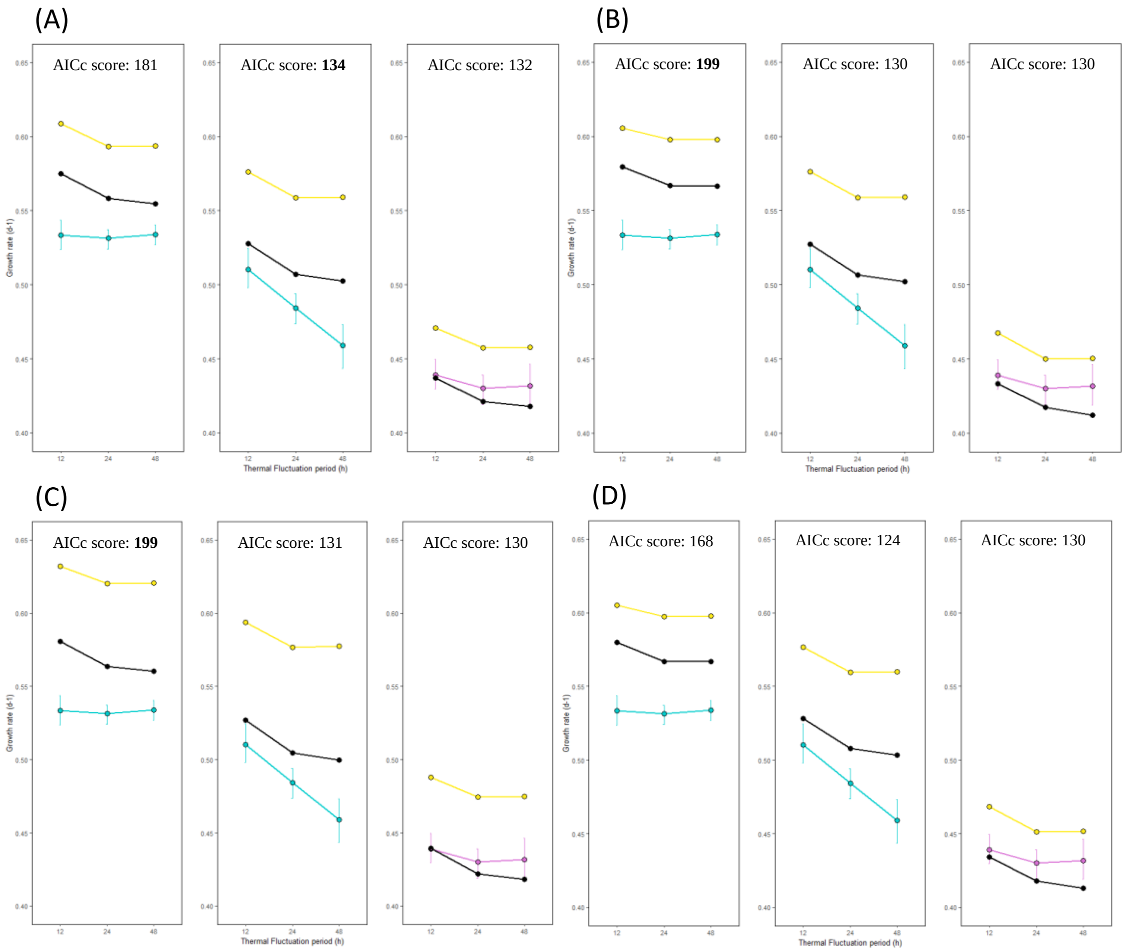

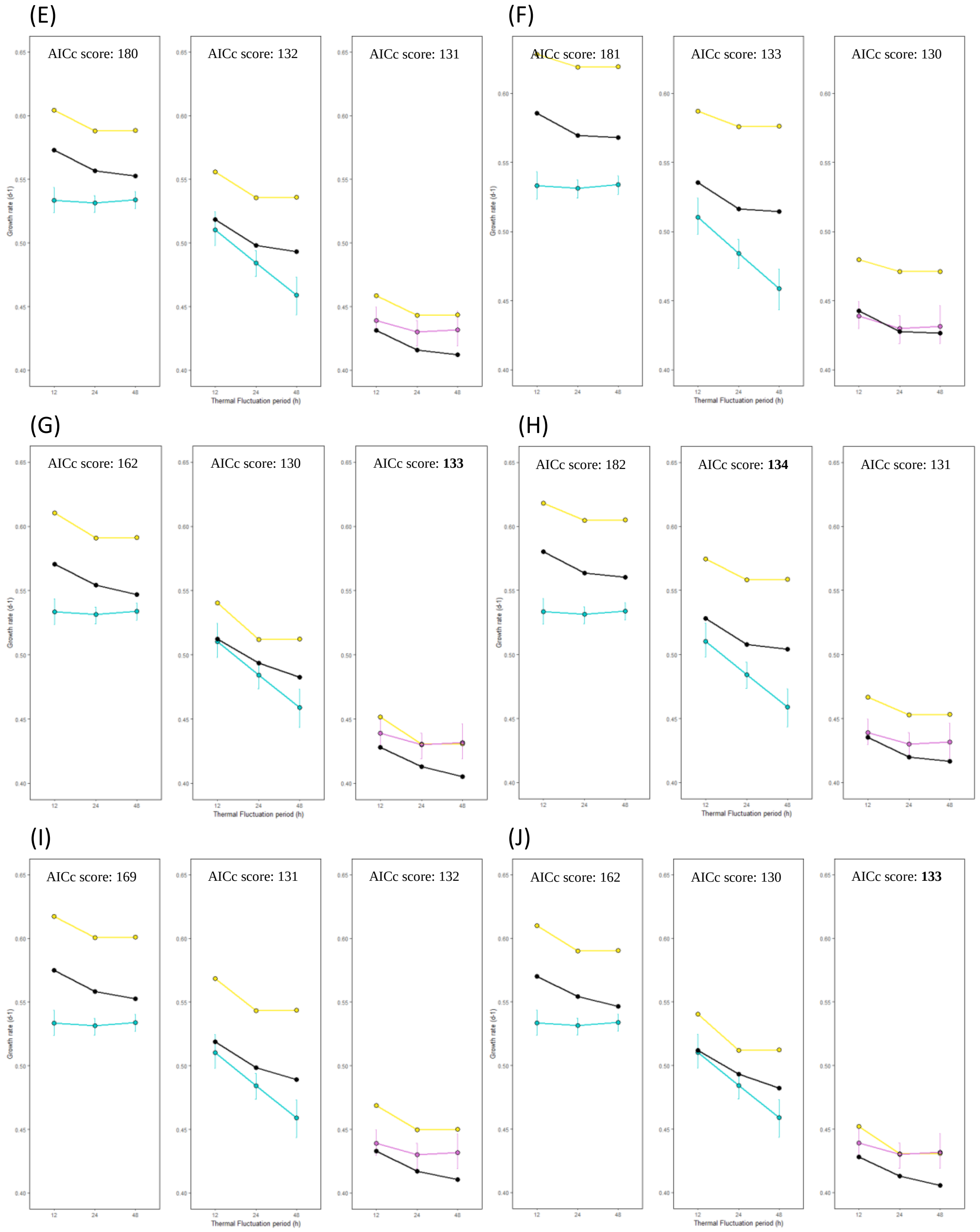
(E)
AICc score: 180
AICc score: 132
AICc score: 131
(F)
AICc score: 181
AICc score: 133
AICc score: 130
(G)
AICc score: 162
AICc score: 130
AICc score: **133**
(H)
AICc score: 182
AICc score: **134**
AICc score: 131
(I)
AICc score: 169
AICc score: 131
AICc score: 132
(J)
AICc score: 162
AICc score: 130
AICc score: **133**
Growth rate (d-1)
Thermal Fluctuation period (h)
12
24
48
0.40
0.45
0.50
0.55
0.60
0.65
