## Supplementary material for "Diet quality alters the consequences of gradual phenotypic plasticity for individual performance": Data and Script: script.R

```
library(boot)
library(car)
library(rTPC)
library(nls.multstart)
library(broom)
library(tidyverse)
library(patchwork)
library(minpack.lm)
library(rstatix)
library(rcompanion)
library(readxl)

setwd("C:.../Supplementary_data/data")
data.0<- read.table("growth_rate_constant.txt", sep="\t", dec=",", header=TRUE)

data.0$M<-data.0$M*1000####mass in mg
mi=8.8##############mean initial mass
data.0$rate<-(log(data.0$M)-log(mi))/(data.0$time*0.0416667)

label_facets_num <- function(string){
  len <- length(string)
  string = paste('(', 1:len, ') ', string, sep = '')
  return(string)
}

data.0<-as_tibble(data.0)%>% 
  dplyr::mutate(temp = as.numeric(temperature))%>%
  dplyr::filter(rate>0)%>%
  dplyr::select(treatement,temp,rate)


nls_multstart_progress <- function(formula, data = parent.frame(), iter, start_lower, 
                                   start_upper, supp_errors = c("Y", "N"), convergence_count = 100, 
                                   control, modelweights, ...){
  if(!is.null(pb)){
    pb$tick()
  }
  nls_multstart(formula = formula, data = data, iter = iter, start_lower = start_lower, 
                start_upper = start_upper, supp_errors = supp_errors, convergence_count = convergence_count, 
                control = control, modelweights = modelweights, ...)
}

# start progress bar and estimate time it will take
number_of_models <- 1
number_of_curves <- length(unique(data.0$treatement))

# setup progress bar
pb <- progress::progress_bar$new(total = number_of_curves*number_of_models,
                                 clear = FALSE,
                                 format ="[:bar] :percent :elapsedfull")


d_fits <- nest(data.0, data = c(temp, rate)) %>%
  mutate(sharpeschoolhigh = map(data, ~nls_multstart(rate~sharpeschoolhigh_1981(temp = temp, r_tref,e,eh,th, tref = 15),
                                                     data = .x,
                                                     iter = c(4,4,4,4),
                                                     start_lower = get_start_vals(.x$temp, .x$rate, model_name = 'sharpeschoolhigh_1981') - 10,
                                                     start_upper = get_start_vals(.x$temp, .x$rate, model_name = 'sharpeschoolhigh_1981') + 10,
                                                     lower = get_lower_lims(.x$temp, .x$rate, model_name = 'sharpeschoolhigh_1981'),
                                                     upper = get_upper_lims(.x$temp, .x$rate, model_name = 'sharpeschoolhigh_1981'),
                                                     supp_errors = 'Y',
                                                     convergence_count = FALSE)))

#d_preds <- mutate(d_fits, new_data = map(data, ~tibble(temp = seq(min(.x$temp), max(.x$temp), length.out = 100)))) %>%
d_preds <- mutate(d_fits, new_data = map(data, ~tibble(temp = seq(16, 33, length.out = 100)))) %>%
  # get rid of original data column
  # select(., -data) %>%
  # stack models into a single column, with an id column for model_name
  pivot_longer(., names_to = 'model_name', values_to = 'fit', c(sharpeschoolhigh))%>%
  # this uses both fit and new_data list columns
  mutate(preds = map2(fit, new_data, ~augment(.x, newdata = .y))) %>%
  # select only the columns we want to keep
  # unlist the preds list column
  unnest(preds)


# get coefs
d_fits <- mutate(d_fits, coefs = map(sharpeschoolhigh, coef))


d_fits <- mutate(d_fits, nls_fit = map2(data, coefs, ~nlsLM(rate~sharpeschoolhigh_1981(temp = temp, r_tref,e,eh,th, tref = 15),
                                                            data = .x,
                                                            start = .y,
                                                            lower = get_lower_lims(.x$temp, .x$rate, model_name = 'sharpeschoolhigh_1981'),
                                                            upper = get_upper_lims(.x$temp, .x$rate, model_name = 'sharpeschoolhigh_1981'))))


# create empty list column
d_fits <- mutate(d_fits, bootstrap = list(rep(NA, n())))

# run for loop to bootstrap each refitted model
for(i in 1:nrow(d_fits)){
  temp_data <- d_fits$data[[i]]
  temp_fit <- nlsLM(rate ~ sharpeschoolhigh_1981(temp = temp, r_tref,e,eh,th, tref = 15),
                    data = temp_data,
                    start = d_fits$coefs[[i]])
  boot <- Boot(temp_fit, method = 'residual', R=1000)
  d_fits$bootstrap[[i]] <- boot
  rm(list = c('temp_fit', 'temp_data', 'boot'))
}


# get the raw values of each bootstrap
d_fits <- mutate(d_fits, output_boot = map(bootstrap, function(x) x$t))

# calculate predictions with a gnarly written function

d_fits <- mutate(d_fits, preds = map2(output_boot, data, function(x, y){
  temp <- as.data.frame(x) %>%
    drop_na() %>%
    mutate(iter = 1:n()) %>%
    group_by_all() %>%
    do(data.frame(temp = seq(min(y$temp), max(y$temp), length.out = 100))) %>%
    ungroup() %>%
    mutate(pred = sharpeschoolhigh_1981(temp = temp, r_tref,e,eh,th, tref = 15))
  return(temp)
}))


data1<- read.table("mean_growth_rate_fluctuation.txt", sep="\t", dec=",", header=TRUE)

data1<-as_tibble(data1)%>% 
  dplyr::mutate(temp=as.numeric(temp))%>%
  dplyr::mutate(g=as.numeric(g))%>%
  dplyr::mutate(lower=as.numeric(lower))%>%
  dplyr::mutate(upper=as.numeric(upper))%>%
  dplyr::select(treatement,type,g,lower,upper,temp)
data1$low<-data1$g-data1$lower


data4<- read_excel("C:....../Supplementary_data/data/growth_rate_fluctuation.xlsx")
data4<-as_tibble(data4)%>% 
  dplyr::mutate(g = as.numeric(g))%>%
  dplyr::filter(Time==150)%>%
  dplyr::select(Treatment,Type,g)


data.0$Treatment<-'CT'
data.0<-as_tibble(data.0)%>% 
  dplyr::mutate(g = as.numeric(rate))%>%
  dplyr::mutate(Type = treatement)%>%
  dplyr::filter(temp==23.80)%>%
  dplyr::select(Treatment,Type,g)

data <- rbind(data4,data.0) 

data=data %>% mutate(treat=str_c(data$Treatment,data$Type,sep="-"))
# cr?ation de la fonction moyenne pour la fonction boot
moyenne<-function(data,indice)
{
  data.star <- data[indice]
  moy <- mean(data.star,na.rm=TRUE)
}

#permet de fixer la graine des tirages al?atoires et donc d'obtenir des r?sultats toujours identiques
set.seed(1234)

# realisation des ?chantillons bootstrap et estimation des moyennes
v.treat<- levels(as.factor(data$treat))
v.1.0 <- NULL
v.2.0 <- NULL
v.3.0 <- NULL

for (i in (1:length(v.treat))) {
  data2<- subset(data, treat == v.treat[i])  
  b1=boot(data2$g, statistic= moyenne, R=1000)
  
  ci=boot.ci(b1,type="perc")
  v<-c(ci[["percent"]])
  v.1<-v[4]
  v.1.0<-c(v.1.0,v.1)
  v.2<-v[5]
  v.2.0<-c(v.2.0,v.2)
  v.3<-mean(data2$g)
  v.3.0<-c(v.3.0,v.3)
  
}
tab.prms <- data.frame(v.treat, v.1.0,v.2.0,v.3.0)
tab.prms ##############g et indice 95
colnames(tab.prms)[colnames(tab.prms) == 'v.treat'] <- 'treat'
data.1=data1 %>% mutate(treat=str_c(data1$type,data1$treatement,sep="-"))
data.4<-left_join(data.1, tab.prms, by ='treat')

data.4<-as_tibble(data.4)%>% 
  dplyr::mutate(rate = as.numeric(v.3.0))%>%
  dplyr::mutate(lower = as.numeric(v.1.0))%>%
  dplyr::mutate(upper = as.numeric(v.2.0))%>%
  dplyr::select(treatement,type,rate,temp,lower,upper)

colnames(data.4)[colnames(data.4) == 'rate'] <- 'g'

k<-8.62e-05

data.F12<- read_excel("C:..../Supplementary_data/data/F12.xlsx")
data.F24<- read_excel("C:..../Supplementary_data/data/F24.xlsx")
data.F48<- read_excel("C:..../Supplementary_data/data/F48.xlsx")
###########################
Treat<-c(rep("F12", length(data.F12$hours)),rep("F24", length(data.F24$hours)),
         rep("F48", length(data.F48$hours)))


A<-ggplot(data.F48,aes(x=hours,y=Temp, color="chartreuse4")) + 
  geom_line(show.legend = FALSE, color="chartreuse4")+ylim(17,30)+  theme_classic()+
  geom_hline(yintercept=22.68, linetype="dashed", color = "chartreuse4")+
  
  theme(axis.text.x = element_text(face="bold",size=10), axis.text.y = element_text(face="bold",size=10),axis.title.x = element_text(size=15),axis.title.y = element_text(size=15))+
  labs(title = "",
       subtitle = "",
       x = "Hours of experiment (h)", y = "",
       tag = "C")

B<-ggplot(data.F24,aes(x=hours,y=Temp, color="turquoise4")) + 
  geom_line(show.legend = FALSE, color="turquoise4")+ylim(17,30)+  theme_classic()+
  geom_hline(yintercept=22.67, linetype="dashed", color = "turquoise4")+
  
  theme(axis.text.x = element_text(face="bold",size=10), axis.text.y = element_text(face="bold",size=10),axis.title.x = element_text(size=15),axis.title.y = element_text(size=15))+
  labs(title = "",
       subtitle = "",
       x = "", y = "Temperature (?C)",
       tag = "B")


C<-ggplot(data.F12,aes(x=hours,y=Temp, color="goldenrod2")) + 
  geom_line(show.legend = FALSE, color="goldenrod2")+ylim(17,30)+  theme_classic()+
  geom_hline(yintercept=23.53, linetype="dashed", color = "goldenrod2")+
  
  theme(axis.text.x = element_text(face="bold",size=10), axis.text.y = element_text(face="bold",size=10),axis.title.x = element_text(size=15),axis.title.y = element_text(size=15))+
  labs(title = "",
       subtitle = "",
       x = "", y = "",
       tag = "A")


library("cowplot")
plot_grid(C,B,A ,ncol = 1, nrow = 3)
#############Y##########

#sharpeschoolhigh_1981
rate <-function(x){
  r_tref*exp((e/k)*(1/tref - 1/(x + 273.15))) / 
    (1 + exp((eh/k)*(1/(th + 273.15) - 1/(x + 273.15))))
}

library(MESS)#####AUC


v.rate.0 <- NULL
v.treat.0 <- NULL
v.Temp.0 <- NULL


v.int6.0<- NULL
###########i=food, j=repbootsrap########
for (i in c(1:3)){
  v.2<-i
  for(j in c(1:1000)){
    
    r_tref<-  d_fits$output_boot[[i]][j,1]
    e<-  d_fits$output_boot[[i]][j,2]
    eh<-  d_fits$output_boot[[i]][j,3]
    th<-  d_fits$output_boot[[i]][j,4]
    
    
    
    
    v.rate<- r_tref*exp((e/k)*(1/288.15 - 1/(data.F12$Temp + 273.15))) / 
      (1 + exp((eh/k)*(1/(th + 273.15) - 1/(data.F12$Temp + 273.15))))
    
    v.1<-mean(v.rate)
    v.4<-mean(data.F12$Temp)  
    v.rate.0 <- c(v.rate.0, v.1 )
    v.Temp.0 <- c(v.Temp.0, v.4 )
    v.treat.0<-c(v.treat.0,v.2)
    v.6<-auc(x=seq(1,length(v.rate),1),y=v.rate)/length(v.rate)
    v.int6.0 <- c(v.int6.0,v.6)     
    
  }
}


tab.prms.rate <- data.frame(v.rate.0, v.Temp.0,v.treat.0,v.int6.0) 

tab.prms.rate$v.treat.0[tab.prms.rate$v.treat.0==1]<-'Y'
tab.prms.rate$v.treat.0[tab.prms.rate$v.treat.0==2]<-'C'
tab.prms.rate$v.treat.0[tab.prms.rate$v.treat.0==3]<-'S'
tab.prms.rate$type<-'F12'

tab.prms.rate<-as_tibble(tab.prms.rate)%>% 
  dplyr::mutate(temp=as.numeric(v.Temp.0))%>%
  dplyr::mutate(g=as.numeric(v.int6.0))%>%
  dplyr::mutate(treatement=v.treat.0)%>%
  dplyr::select(treatement,g,temp,type)##########mean temperature = 23.5317

v.rate.0 <- NULL
v.treat.0 <- NULL
v.Temp.0 <- NULL
###########i=food, j=repbootsrap########
for (i in c(1:3)){
  v.2<-i
  for(j in c(1:1000)){
    
    r_tref<-  d_fits$output_boot[[i]][j,1]
    e<-  d_fits$output_boot[[i]][j,2]
    eh<-  d_fits$output_boot[[i]][j,3]
    th<-  d_fits$output_boot[[i]][j,4]
    
    
    
    
    v.rate<- r_tref*exp((e/k)*(1/288.15 - 1/(23.5317 + 273.15))) / 
      (1 + exp((eh/k)*(1/(th + 273.15) - 1/(23.5317 + 273.15))))
    
    v.1<-v.rate
    v.4<-23.5317
    v.rate.0 <- c(v.rate.0, v.1 )
    v.Temp.0 <- c(v.Temp.0, v.4 )
    v.treat.0<-c(v.treat.0,v.2)
    
  }
}


tab.prms.rate.ctt <- data.frame(v.rate.0, v.Temp.0,v.treat.0) 

tab.prms.rate.ctt$v.treat.0[tab.prms.rate.ctt$v.treat.0==1]<-'Y'
tab.prms.rate.ctt$v.treat.0[tab.prms.rate.ctt$v.treat.0==2]<-'C'
tab.prms.rate.ctt$v.treat.0[tab.prms.rate.ctt$v.treat.0==3]<-'S'
tab.prms.rate.ctt$type<-'F12'

tab.prms.rate.ctt<-as_tibble(tab.prms.rate.ctt)%>% 
  dplyr::mutate(temp=as.numeric(v.Temp.0))%>%
  dplyr::mutate(g=as.numeric(v.rate.0))%>%
  dplyr::mutate(treatement=v.treat.0)%>%
  dplyr::select(treatement,g,temp,type)##########mean temperature = 23.5317

#############period 24##########


v.rate.0 <- NULL
v.treat.0 <- NULL
v.Temp.0 <- NULL

v.int6.0<- NULL


for (i in c(1:3)){
  v.2<-i
  for(j in c(1:1000)){
    
    r_tref<-  d_fits$output_boot[[i]][j,1]
    e<-  d_fits$output_boot[[i]][j,2]
    eh<-  d_fits$output_boot[[i]][j,3]
    th<-  d_fits$output_boot[[i]][j,4]
    
    
    
    
    v.rate<- r_tref*exp((e/k)*(1/288.15 - 1/(data.F24$Temp + 273.15))) / 
      (1 + exp((eh/k)*(1/(th + 273.15) - 1/(data.F24$Temp + 273.15))))
    
    v.1<-mean(v.rate)
    v.4<-mean(data.F24$Temp)  
    v.rate.0 <- c(v.rate.0, v.1 )
    v.Temp.0 <- c(v.Temp.0, v.4 )
    v.treat.0<-c(v.treat.0,v.2)
    
    v.6<-auc(x=seq(1,length(v.rate),1),y=v.rate)/length(v.rate)
    v.int6.0 <- c(v.int6.0,v.6)       
    
  }
}


tab.prms.rate.24 <- data.frame(v.rate.0, v.Temp.0,v.treat.0,v.int6.0) 

tab.prms.rate.24$v.treat.0[tab.prms.rate.24$v.treat.0==1]<-'Y'
tab.prms.rate.24$v.treat.0[tab.prms.rate.24$v.treat.0==2]<-'C'
tab.prms.rate.24$v.treat.0[tab.prms.rate.24$v.treat.0==3]<-'S'
tab.prms.rate.24$type<-'F24'

tab.prms.rate.24<-as_tibble(tab.prms.rate.24)%>% 
  dplyr::mutate(temp=as.numeric(v.Temp.0))%>%
  dplyr::mutate(g=as.numeric(v.int6.0))%>%
  dplyr::mutate(treatement=v.treat.0)%>%
  dplyr::select(treatement,g,temp,type)###########mean temperature =22.66641
v.rate.0 <- NULL
v.treat.0 <- NULL
v.Temp.0 <- NULL
###########i=food, j=repbootsrap########
for (i in c(1:3)){
  v.2<-i
  for(j in c(1:1000)){
    
    r_tref<-  d_fits$output_boot[[i]][j,1]
    e<-  d_fits$output_boot[[i]][j,2]
    eh<-  d_fits$output_boot[[i]][j,3]
    th<-  d_fits$output_boot[[i]][j,4]
    
    
    
    
    v.rate<- r_tref*exp((e/k)*(1/288.15 - 1/(22.66641 + 273.15))) / 
      (1 + exp((eh/k)*(1/(th + 273.15) - 1/(22.66641 + 273.15))))
    
    v.1<-v.rate
    v.4<-22.66641
    v.rate.0 <- c(v.rate.0, v.1 )
    v.Temp.0 <- c(v.Temp.0, v.4 )
    v.treat.0<-c(v.treat.0,v.2)
    
  }
}


tab.prms.rate.24.ctt <- data.frame(v.rate.0, v.Temp.0,v.treat.0) 

tab.prms.rate.24.ctt$v.treat.0[tab.prms.rate.24.ctt$v.treat.0==1]<-'Y'
tab.prms.rate.24.ctt$v.treat.0[tab.prms.rate.24.ctt$v.treat.0==2]<-'C'
tab.prms.rate.24.ctt$v.treat.0[tab.prms.rate.24.ctt$v.treat.0==3]<-'S'
tab.prms.rate.24.ctt$type<-'F24'

tab.prms.rate.24.ctt<-as_tibble(tab.prms.rate.24.ctt)%>% 
  dplyr::mutate(temp=as.numeric(v.Temp.0))%>%
  dplyr::mutate(g=as.numeric(v.rate.0))%>%
  dplyr::mutate(treatement=v.treat.0)%>%
  dplyr::select(treatement,g,temp,type)

#####################period48###########

v.rate.0 <- NULL
v.treat.0 <- NULL
v.Temp.0 <- NULL


v.int6.0<- NULL

for (i in c(1:3)){
  v.2<-i
  for(j in c(1:1000)){
    
    r_tref<-  d_fits$output_boot[[i]][j,1]
    e<-  d_fits$output_boot[[i]][j,2]
    eh<-  d_fits$output_boot[[i]][j,3]
    th<-  d_fits$output_boot[[i]][j,4]
    
    
    
    
    v.rate<- r_tref*exp((e/k)*(1/288.15 - 1/(data.F48$Temp + 273.15))) / 
      (1 + exp((eh/k)*(1/(th + 273.15) - 1/(data.F48$Temp + 273.15))))
    
    v.1<-mean(v.rate)
    v.4<-mean(data.F48$Temp)  
    v.rate.0 <- c(v.rate.0, v.1 )
    v.Temp.0 <- c(v.Temp.0, v.4 )
    v.treat.0<-c(v.treat.0,v.2)
    v.6<-auc(x=seq(1,length(v.rate),1),y=v.rate)/length(v.rate)
    v.int6.0 <- c(v.int6.0,v.6) 
  }
}


tab.prms.rate.48 <- data.frame(v.rate.0, v.Temp.0,v.treat.0,v.int6.0) 

tab.prms.rate.48$v.treat.0[tab.prms.rate.48$v.treat.0==1]<-'Y'
tab.prms.rate.48$v.treat.0[tab.prms.rate.48$v.treat.0==2]<-'C'
tab.prms.rate.48$v.treat.0[tab.prms.rate.48$v.treat.0==3]<-'S'
tab.prms.rate.48$type<-'F48'


tab.prms.rate.48<-as_tibble(tab.prms.rate.48)%>% 
  dplyr::mutate(temp=as.numeric(v.Temp.0))%>%
  dplyr::mutate(g=as.numeric(v.int6.0))%>%
  dplyr::mutate(treatement=v.treat.0)%>%
  dplyr::select(treatement,g,temp,type)####mean temperature =22.68369
v.rate.0 <- NULL
v.treat.0 <- NULL
v.Temp.0 <- NULL
###########i=food, j=repbootsrap########
for (i in c(1:3)){
  v.2<-i
  for(j in c(1:1000)){
    
    r_tref<-  d_fits$output_boot[[i]][j,1]
    e<-  d_fits$output_boot[[i]][j,2]
    eh<-  d_fits$output_boot[[i]][j,3]
    th<-  d_fits$output_boot[[i]][j,4]
    
    
    
    
    v.rate<- r_tref*exp((e/k)*(1/288.15 - 1/(22.68369 + 273.15))) / 
      (1 + exp((eh/k)*(1/(th + 273.15) - 1/(22.68369 + 273.15))))
    
    v.1<-v.rate
    v.4<-22.68369
    v.rate.0 <- c(v.rate.0, v.1 )
    v.Temp.0 <- c(v.Temp.0, v.4 )
    v.treat.0<-c(v.treat.0,v.2)
    
  }
}


tab.prms.rate.48.ctt <- data.frame(v.rate.0, v.Temp.0,v.treat.0) 

tab.prms.rate.48.ctt$v.treat.0[tab.prms.rate.48.ctt$v.treat.0==1]<-'Y'
tab.prms.rate.48.ctt$v.treat.0[tab.prms.rate.48.ctt$v.treat.0==2]<-'C'
tab.prms.rate.48.ctt$v.treat.0[tab.prms.rate.48.ctt$v.treat.0==3]<-'S'
tab.prms.rate.48.ctt$type<-'F48'

tab.prms.rate.48.ctt<-as_tibble(tab.prms.rate.48.ctt)%>% 
  dplyr::mutate(temp=as.numeric(v.Temp.0))%>%
  dplyr::mutate(g=as.numeric(v.rate.0))%>%
  dplyr::mutate(treatement=v.treat.0)%>%
  dplyr::select(treatement,g,temp,type)


data_freq<-rbind(tab.prms.rate,tab.prms.rate.24,tab.prms.rate.48)
data_freq_ctt<-rbind(tab.prms.rate.ctt,tab.prms.rate.24.ctt,tab.prms.rate.48.ctt)

pd=position_dodge(0.3)


dataf<- read_excel("C:....../Supplementary_data/data/growth_rate_fluctuation.xlsx")

dataf<-dataf[-which(dataf$Treatment =='cst'),]
dataf= dataf %>% filter(Time=="150")
dataf<-as_tibble(dataf)%>% 
  dplyr::mutate(g=as.numeric(g))%>%
  dplyr::mutate(type=Treatment)%>%
  dplyr::mutate(treatement=Type)%>%
  dplyr::select(treatement,g,type)

data_freq<-as_tibble(data_freq)%>% 
  dplyr::select(treatement,g,type)
data_freq_ctt<-as_tibble(data_freq_ctt)%>% 
  dplyr::select(treatement,g,type)

dataf$class<- 'obs'
data_freq$class<-'pred'
data_freq_ctt$class<-'ctt'
data<-rbind(dataf,data_freq,data_freq_ctt)

data=data %>% mutate(treat=str_c(data$treatement,data$type,data$class,sep="-"))
# cr?ation de la fonction moyenne pour la fonction boot
moyenne<-function(data,indice)
{
  data.star <- data[indice]
  moy <- mean(data.star,na.rm=TRUE)
}

#permet de fixer la graine des tirages al?atoires et donc d'obtenir des r?sultats toujours identiques
set.seed(1234)

# realisation des ?chantillons bootstrap et estimation des moyennes
v.treat<- levels(as.factor(data$treat))
v.1.0 <- NULL
v.2.0 <- NULL
v.3.0 <- NULL
v.4.0 <- NULL
v.5.0 <- NULL
v.6.0 <- NULL
for (i in (1:length(v.treat))) {
  data2<- subset(data, treat == v.treat[i])  
  b1=boot(data2$g, statistic= moyenne, R=1000)
  
  ci=boot.ci(b1,type="perc")
  v<-c(ci[["percent"]])
  v.1<-v[4]
  v.1.0<-c(v.1.0,v.1)
  v.2<-v[5]
  v.2.0<-c(v.2.0,v.2)
  v.3<-mean(data2$g)
  v.3.0<-c(v.3.0,v.3)
  v.4<-first(data2$treatement)
  v.4.0<-c(v.4.0,v.4)
  v.5<-first(data2$type)
  v.5.0<-c(v.5.0,v.5)
  v.6<-first(data2$class)
  v.6.0<-c(v.6.0,v.6)
}
tab.prms <- data.frame(v.treat, v.1.0,v.2.0,v.3.0,v.4.0,v.5.0,v.6.0)
tab.prms ##############g et indice 95
colnames(tab.prms)[colnames(tab.prms) == 'v.treat'] <- 'treat'


data<-as_tibble(tab.prms)%>% 
  dplyr::mutate(rate = as.numeric(v.3.0))%>%
  dplyr::mutate(lower = as.numeric(v.1.0))%>%
  dplyr::mutate(upper = as.numeric(v.2.0))%>%
  dplyr::mutate(treatement = v.4.0)%>%
  dplyr::mutate(type = v.5.0)%>%
  dplyr::mutate(class = v.6.0)%>%
  dplyr::select(treatement,type,rate,class,lower,upper)

colnames(data)[colnames(data) == 'rate'] <- 'g'


pd=position_dodge(0) 


data.Y<-as_tibble(data)%>% 
  dplyr::filter(treatement=="Y")
data.Y$type[data.Y$type=='F12']<-12
data.Y$type[data.Y$type=='F24']<-24
data.Y$type[data.Y$type=='F48']<-48
a<-ggplot(data.Y,aes(x=type, y=g,group = class,col=class),show.legend = FALSE) +
  geom_errorbar(aes(x=type,y=g,ymin=lower,ymax=upper),data.Y,fill=class, width=.1, size=0.15,show.legend = FALSE,  position=pd)+
  geom_point(size=3,show.legend = FALSE)+
  geom_line(size=1,show.legend = FALSE)+
  geom_point(shape = 1,size = 3,colour = "black")+
  theme(  panel.background = element_blank(),
          panel.grid.major = element_blank(), 
          panel.grid.minor = element_blank(),
          panel.border = element_rect(colour = "black",fill=NA))+
  scale_color_manual(breaks = c("obs", "pred","ctt"),values=c("cyan3","black","#ffec1b"),
                     labels=c("Observation","Prediction","Constant"))+
  
  labs(x = '',
       y = 'Growth rate (d-1)',title='')+
  theme(axis.text.x = element_text(size=10),axis.text.y = element_text(size=10),
        axis.title.x = element_text(size=14),axis.title.y = element_text(size=14))+ylim(0.40,0.63)


data.C<-as_tibble(data)%>% 
  dplyr::filter(treatement=="C")
data.C$type[data.C$type=='F12']<-12
data.C$type[data.C$type=='F24']<-24
data.C$type[data.C$type=='F48']<-48
b<-ggplot(data.C,aes(x=type, y=g,group = class,col=class),show.legend = FALSE) + 
  geom_errorbar(aes(x=type,y=g,ymin=lower,ymax=upper),data.C,fill=class, width=.1, size=0.15,show.legend = FALSE,  position=pd)+
  geom_point(size=3,show.legend = FALSE)+
  geom_line(size=1,show.legend = FALSE)+
  geom_point(shape = 1,size = 3,colour = "black")+
  theme(  panel.background = element_blank(),
          panel.grid.major = element_blank(), 
          panel.grid.minor = element_blank(),
          panel.border = element_rect(colour = "black",fill=NA))+
  scale_color_manual(breaks = c("obs", "pred","ctt"),values=c("cyan3","black","#ffec1b"),
                     labels=c("Observation","Prediction","Constant"))+
  
  labs(x = 'Thermal Fluctuation period (h)',
       y = '',title='')+
  theme(axis.text.x = element_text(size=10),axis.text.y = element_text(size=10),
        axis.title.x = element_text(size=14),axis.title.y = element_text(size=14))+ylim(0.40,0.63)

data.S<-as_tibble(data)%>% 
  dplyr::filter(treatement=="S")
data.S$type[data.S$type=='F12']<-12
data.S$type[data.S$type=='F24']<-24
data.S$type[data.S$type=='F48']<-48
c<-ggplot(data.S,aes(x=type, y=g,group = class,col=class),show.legend = FALSE) +
  geom_errorbar(aes(x=type,y=g,ymin=lower,ymax=upper),data.S,fill=class, width=.1, size=0.15,show.legend = FALSE,  position=pd)+
  geom_point(size=3,show.legend = FALSE)+
  geom_line(size=1,show.legend = FALSE)+
  geom_point(shape = 1,size = 3,colour = "black")+
  theme(  panel.background = element_blank(),
          panel.grid.major = element_blank(), 
          panel.grid.minor = element_blank(),
          panel.border = element_rect(colour = "black",fill=NA))+
  scale_color_manual(breaks = c("obs", "pred","ctt"),values=c("orchid","black","#ffec1b"),
                     labels=c("Observation","Prediction","Constant"))+
  labs(x = '',
       y = '',title='')+
  theme(axis.text.x = element_text(size=10),axis.text.y = element_text(size=10),
        axis.title.x = element_text(size=14),axis.title.y = element_text(size=14))+ylim(0.40,0.63)


library("cowplot")
plot_grid(a,b,c ,ncol = 3, nrow = 1)


####################################


data.mean=data %>% mutate(treat=str_c(data$type,data$class,data$treatement,sep="-"))
data.mean=data.mean %>% group_by(treat) %>% summarise(treatement=first(treatement),class= first(class),type=first(type),
                                                      g=mean(g))

library(MKinfer)
pred.C.fast.f<-data_freq %>% filter(type=="F12" & treatement== "C")
obs.C.fast.f <-dataf %>% filter(type=="F12" & treatement== "C")
ctt.C.fast.f<-data_freq_ctt %>% filter(type=="F12" & treatement== "C")

pred.C.mid.f<-data_freq %>% filter(type=="F24" & treatement== "C")
obs.C.mid.f <-dataf %>% filter(type=="F24" & treatement== "C")
ctt.C.mid.f<-data_freq_ctt %>% filter(type=="F24" & treatement== "C")

pred.C.slow.f<-data_freq %>% filter(type=="F48" & treatement== "C")
obs.C.slow.f <-dataf %>% filter(type=="F48" & treatement== "C")
ctt.C.slow.f<-data_freq_ctt %>% filter(type=="F48" & treatement== "C")

pred.Y.fast.f<-data_freq %>% filter(type=="F12" & treatement== "Y")
obs.Y.fast.f <-dataf %>% filter(type=="F12" & treatement== "Y")
ctt.Y.fast.f<-data_freq_ctt %>% filter(type=="F12" & treatement== "Y")

pred.Y.mid.f<-data_freq %>% filter(type=="F24" & treatement== "Y")
obs.Y.mid.f <-dataf %>% filter(type=="F24" & treatement== "Y")
ctt.Y.mid.f<-data_freq_ctt %>% filter(type=="F24" & treatement== "Y")

pred.Y.slow.f<-data_freq %>% filter(type=="F48" & treatement== "Y")
obs.Y.slow.f <-dataf %>% filter(type=="F48" & treatement== "Y")
ctt.Y.slow.f<-data_freq_ctt %>% filter(type=="F48" & treatement== "Y")

pred.S.fast.f<-data_freq %>% filter(type=="F12" & treatement== "S")
obs.S.fast.f <-dataf %>% filter(type=="F12" & treatement== "S")
ctt.S.fast.f<-data_freq_ctt %>% filter(type=="F12" & treatement== "S")

pred.S.mid.f<-data_freq %>% filter(type=="F24" & treatement== "S")
obs.S.mid.f <-dataf %>% filter(type=="F24" & treatement== "S")
ctt.S.mid.f<-data_freq_ctt %>% filter(type=="F24" & treatement== "S")

pred.S.slow.f<-data_freq %>% filter(type=="F48" & treatement== "S")
obs.S.slow.f <-dataf %>% filter(type=="F48" & treatement== "S")
ctt.S.slow.f<-data_freq_ctt %>% filter(type=="F48" & treatement== "S")

boot.t.test(x=pred.Y.slow.f$g, y=obs.Y.slow.f$g)
boot.t.test(x=pred.Y.mid.f$g, y=obs.Y.mid.f$g)
boot.t.test(x=pred.Y.fast.f$g, y=obs.Y.fast.f$g)

boot.t.test(x=pred.C.slow.f$g, y=obs.C.slow.f$g)
boot.t.test(x=pred.C.mid.f$g, y=obs.C.mid.f$g)
boot.t.test(x=pred.C.fast.f$g, y=obs.C.fast.f$g)

boot.t.test(x=pred.S.slow.f$g, y=obs.S.slow.f$g)
boot.t.test(x=pred.S.mid.f$g, y=obs.S.mid.f$g)
boot.t.test(x=pred.S.fast.f$g, y=obs.S.fast.f$g)


boot.t.test(x=ctt.Y.slow.f$g, y=obs.Y.slow.f$g)
boot.t.test(x=ctt.Y.mid.f$g, y=obs.Y.mid.f$g)
boot.t.test(x=ctt.Y.fast.f$g, y=obs.Y.fast.f$g)

boot.t.test(x=ctt.C.slow.f$g, y=obs.C.slow.f$g)
boot.t.test(x=ctt.C.mid.f$g, y=obs.C.mid.f$g)
boot.t.test(x=ctt.C.fast.f$g, y=obs.C.fast.f$g)

boot.t.test(x=ctt.S.slow.f$g, y=obs.S.slow.f$g)
boot.t.test(x=ctt.S.mid.f$g, y=obs.S.mid.f$g)
boot.t.test(x=ctt.S.fast.f$g, y=obs.S.fast.f$g)
###############


#############anova#############

library(performance)
library(emmeans)
TG.aov <- aov(g~treatement*type, 
              contrasts = list(treatement=contr.sum,
                               type=contr.sum),
              data=dataf)
Anova(TG.aov, type=3)

check_normality(TG.aov)
check_homogeneity(TG.aov)

pwc <- dataf%>% 
  group_by(treatement) %>%
  emmeans_test(g ~ type, p.adjust.method = "bonferroni") 
pwc

pwc1 <- dataf%>% 
  group_by(type) %>%
  emmeans_test(g ~ treatement, p.adjust.method = "bonferroni") 
pwc1
###############################

v.rate.0 <- NULL
v.upper.0 <- NULL
v.lower.0 <- NULL
v.Temp.0 <- NULL
v.int6.0<-NULL
#v.int7.0<-NULL

for (i in c(1:116)){
  data2 <- subset(data.F12)
  data2$Temp<-data2$Temp +0.1
  
  v.rate<- 0.3584294*exp((0.5810565/k)*(1/288.15 - 1/(data2$Temp + 273.15))) / 
    (1 + exp((2.5680691/k)*(1/(29.2655841 + 273.15) - 1/(data2$Temp + 273.15))))
  v.lower <-  0.3269024    *exp((0.4172143/k)*(1/288.15 - 1/(data2$Temp + 273.15))) / 
    (1 + exp((2.1363105  /k)*(1/(26.5547148 + 273.15) - 1/(data2$Temp + 273.15))))
  v.upper <-0.3913538*exp((0.8316800/k)*(1/288.15 - 1/(data2$Temp + 273.15))) / 
    (1 + exp((3.1262846/k)*(1/(30.9481058+ 273.15) - 1/(data2$Temp + 273.15))))
  
  v.1<-mean(v.rate)
  v.2<-mean(v.lower)
  v.3<-mean(v.upper)
  v.4<-mean(data2$Temp)
  
  v.rate.0 <- c(v.rate.0, v.1 )
  v.upper.0 <- c(v.upper.0, v.2 )
  v.lower.0 <- c(v.lower.0, v.3 )
  v.Temp.0 <- c(v.Temp.0, v.4 )
  
  v.6<-auc(x=seq(1,length(v.rate),1),y=v.rate)/length(v.rate)
  
  
  v.int6.0 <- c(v.int6.0,v.6) 
  
  data.F12$Temp<-data2$Temp  
  
}

tab.prms.rate.Y.1  <- data.frame(v.rate.0, v.upper.0, v.lower.0,v.Temp.0,v.int6.0)
data.F12<- read_excel("C:....../Supplementary_data/data/F12.xlsx")

v.rate.0 <- NULL
v.upper.0 <- NULL
v.lower.0 <- NULL
v.Temp.0 <- NULL
v.int6.0<-NULL


for (i in c(1:116)){
  data2 <- subset(data.F12)
  data2$Temp<-data2$Temp -0.1
  
  v.rate<- 0.3584294*exp((0.5810565/k)*(1/288.15 - 1/(data2$Temp + 273.15))) / 
    (1 + exp((2.5680691/k)*(1/(29.2655841 + 273.15) - 1/(data2$Temp + 273.15))))
  v.lower <-  0.3269024    *exp((0.4172143/k)*(1/288.15 - 1/(data2$Temp + 273.15))) / 
    (1 + exp((2.1363105  /k)*(1/(26.5547148 + 273.15) - 1/(data2$Temp + 273.15))))
  v.upper <-0.3913538*exp((0.8316800/k)*(1/288.15 - 1/(data2$Temp + 273.15))) / 
    (1 + exp((3.1262846/k)*(1/(30.9481058+ 273.15) - 1/(data2$Temp + 273.15))))
  
  
  v.1<-mean(v.rate)
  v.2<-mean(v.lower)
  v.3<-mean(v.upper)
  v.4<-mean(data2$Temp)
  v.rate.0 <- c(v.rate.0, v.1 )
  v.upper.0 <- c(v.upper.0, v.2 )
  v.lower.0 <- c(v.lower.0, v.3 )
  v.Temp.0 <- c(v.Temp.0, v.4 )
  
  v.6<-auc(x=seq(1,length(v.rate),1),y=v.rate)/length(v.rate)
  v.int6.0 <- c(v.int6.0,v.6) 
  
  
  data.F12$Temp<-data2$Temp  
}

tab.prms.rate.Y.2  <- data.frame(v.rate.0, v.upper.0, v.lower.0,v.Temp.0,v.int6.0)
data.Y<- rbind(tab.prms.rate.Y.1,tab.prms.rate.Y.2)
####################C#######
data.F12<- read_excel("C:....../Supplementary_data/data/F12.xlsx")
v.rate.0 <- NULL
v.upper.0 <- NULL
v.lower.0 <- NULL
v.Temp.0 <- NULL
v.int6.0<-NULL


for (i in c(1:116)){
  data2 <- subset(data.F12)
  data2$Temp<-data2$Temp +0.1
  
  v.rate<-  0.2835495*exp((0.5840116/k)*(1/288.15 - 1/(data2$Temp + 273.15))) / 
    (1 + exp((4.1664022/k)*(1/(29.9199098 + 273.15) - 1/(data2$Temp + 273.15))))
  v.lower <-  0.2471952    *exp((0.4347565  /k)*(1/288.15 - 1/(data2$Temp + 273.15))) / 
    (1 + exp((3.3303797   /k)*(1/(28.8537428  + 273.15) - 1/(data2$Temp+ 273.15))))
  v.upper <- 0.3192488*exp((0.7496537/k)*(1/288.15 - 1/(data2$Temp + 273.15))) / 
    (1 + exp((5.3575241/k)*(1/(30.7212264 + 273.15) - 1/(data2$Temp + 273.15))))
  
  v.1<-mean(v.rate)
  v.2<-mean(v.lower)
  v.3<-mean(v.upper)
  v.4<-mean(data2$Temp)
  v.rate.0 <- c(v.rate.0, v.1 )
  v.upper.0 <- c(v.upper.0, v.2 )
  v.lower.0 <- c(v.lower.0, v.3 )
  v.Temp.0 <- c(v.Temp.0, v.4 )
  v.6<-auc(x=seq(1,length(v.rate),1),y=v.rate)/length(v.rate)
  v.int6.0 <- c(v.int6.0,v.6) 
  
  data.F12$Temp<-data2$Temp  
  
}

tab.prms.rate.C.1  <- data.frame(v.rate.0, v.upper.0, v.lower.0,v.Temp.0,v.int6.0)
data.F12<- read_excel("C:....../Supplementary_data/data/F12.xlsx")

v.rate.0 <- NULL
v.upper.0 <- NULL
v.lower.0 <- NULL
v.Temp.0 <- NULL
v.int6.0<-NULL

for (i in c(1:116)){
  data2 <- subset(data.F12)
  data2$Temp<-data2$Temp -0.1
  
  v.rate<-  0.2835495*exp((0.5840116/k)*(1/288.15 - 1/(data2$Temp + 273.15))) / 
    (1 + exp((4.1664022/k)*(1/(29.9199098 + 273.15) - 1/(data2$Temp + 273.15))))
  v.lower <-  0.2471952    *exp((0.4347565  /k)*(1/288.15 - 1/(data2$Temp + 273.15))) / 
    (1 + exp((3.3303797   /k)*(1/(28.8537428  + 273.15) - 1/(data2$Temp+ 273.15))))
  v.upper <- 0.3192488*exp((0.7496537/k)*(1/288.15 - 1/(data2$Temp + 273.15))) / 
    (1 + exp((5.3575241/k)*(1/(30.7212264 + 273.15) - 1/(data2$Temp + 273.15))))
  
  v.1<-mean(v.rate)
  v.2<-mean(v.lower)
  v.3<-mean(v.upper)
  v.4<-mean(data2$Temp)
  v.rate.0 <- c(v.rate.0, v.1 )
  v.upper.0 <- c(v.upper.0, v.2 )
  v.lower.0 <- c(v.lower.0, v.3 )
  v.Temp.0 <- c(v.Temp.0, v.4 )
  v.6<-auc(x=seq(1,length(v.rate),1),y=v.rate)/length(v.rate)
  v.int6.0 <- c(v.int6.0,v.6) 
  
  data.F12$Temp<-data2$Temp  
}

tab.prms.rate.C.2  <- data.frame(v.rate.0, v.upper.0, v.lower.0,v.Temp.0,v.int6.0)
data.C<- rbind(tab.prms.rate.C.1,tab.prms.rate.C.2)

##################S###############
data.F12<- read_excel("C:....../Supplementary_data/data/F12.xlsx")
v.rate.0 <- NULL
v.upper.0 <- NULL
v.lower.0 <- NULL
v.Temp.0 <- NULL
v.int6.0<-NULL

for (i in c(1:116)){
  data2 <- subset(data.F12)
  data2$Temp<-data2$Temp +0.1
  
  v.rate<-  0.2463239*exp((0.5609465/k)*(1/288.15 - 1/(data2$Temp + 273.15))) / 
    (1 + exp((3.6185556/k)*(1/(30.1368399 + 273.15) - 1/(data2$Temp + 273.15))))
  v.lower <-  0.2059233  *exp((0.3687229  /k)*(1/288.15 - 1/(data2$Temp + 273.15))) / 
    (1 + exp((2.7569751  /k)*(1/(27.8246672 + 273.15) - 1/(data2$Temp + 273.15))))
  v.upper <-0.2863341*exp((0.8269036/k)*(1/288.15 - 1/(data2$Temp + 273.15))) / 
    (1 + exp((5.1204435/k)*(1/(31.4278000 + 273.15) - 1/(data2$Temp + 273.15))))
  
  v.1<-mean(v.rate)
  v.2<-mean(v.lower)
  v.3<-mean(v.upper)
  v.4<-mean(data2$Temp)
  v.rate.0 <- c(v.rate.0, v.1 )
  v.upper.0 <- c(v.upper.0, v.2 )
  v.lower.0 <- c(v.lower.0, v.3 )
  v.Temp.0 <- c(v.Temp.0, v.4 )
  v.6<-auc(x=seq(1,length(v.rate),1),y=v.rate)/length(v.rate)
  v.int6.0 <- c(v.int6.0,v.6) 
  
  data.F12$Temp<-data2$Temp  
  
}
data.F12<- read_excel("C:....../Supplementary_data/data/F12.xlsx")
tab.prms.rate.S.1  <- data.frame(v.rate.0, v.upper.0, v.lower.0,v.Temp.0,v.int6.0)


v.rate.0 <- NULL
v.upper.0 <- NULL
v.lower.0 <- NULL
v.Temp.0 <- NULL
v.int6.0<-NULL


for (i in c(1:116)){
  data2 <- subset(data.F12)
  data2$Temp<-data2$Temp -0.1
  
  v.rate<-  0.2463239*exp((0.5609465/k)*(1/288.15 - 1/(data2$Temp + 273.15))) / 
    (1 + exp((3.6185556/k)*(1/(30.1368399 + 273.15) - 1/(data2$Temp + 273.15))))
  v.lower <-  0.2059233  *exp((0.3687229  /k)*(1/288.15 - 1/(data2$Temp + 273.15))) / 
    (1 + exp((2.7569751  /k)*(1/(27.8246672 + 273.15) - 1/(data2$Temp + 273.15))))
  v.upper <-0.2863341*exp((0.8269036/k)*(1/288.15 - 1/(data2$Temp + 273.15))) / 
    (1 + exp((5.1204435/k)*(1/(31.4278000 + 273.15) - 1/(data2$Temp + 273.15))))
  
  
  v.1<-mean(v.rate)
  v.2<-mean(v.lower)
  v.3<-mean(v.upper)
  v.4<-mean(data2$Temp)
  v.rate.0 <- c(v.rate.0, v.1 )
  v.upper.0 <- c(v.upper.0, v.2 )
  v.lower.0 <- c(v.lower.0, v.3 )
  v.Temp.0 <- c(v.Temp.0, v.4 )
  v.6<-auc(x=seq(1,length(v.rate),1),y=v.rate)/length(v.rate)
  v.int6.0 <- c(v.int6.0,v.6) 
  
  data.F12$Temp<-data2$Temp  
}

tab.prms.rate.S.2  <- data.frame(v.rate.0, v.upper.0, v.lower.0,v.Temp.0,v.int6.0)
data.S<- rbind(tab.prms.rate.S.1,tab.prms.rate.S.2)

###############
data.Y$treatement<-'Y'
data.C$treatement<-'C'
data.S$treatement<-'S'
data.all<-rbind(data.Y,data.C,data.S)

#############
v.rate.0 <- NULL
v.upper.0 <- NULL
v.lower.0 <- NULL
v.Temp.0 <- NULL
v.int6.0<-NULL

data.F24<- read_excel("C:....../Supplementary_data/data/F24.xlsx")
for (i in c(1:116)){
  data2 <- subset(data.F24)
  data2$Temp<-data2$Temp +0.1
  
  v.rate<- 0.3584294*exp((0.5810565/k)*(1/288.15 - 1/(data2$Temp + 273.15))) / 
    (1 + exp((2.5680691/k)*(1/(29.2655841 + 273.15) - 1/(data2$Temp + 273.15))))
  v.lower <-  0.4996351  *exp((0.4164426/k)*(1/288.15 - 1/(data2$Temp + 273.15))) / 
    (1 + exp((2.1743233/k)*(1/(26.8258663 + 273.15) - 1/(data2$Temp + 273.15))))
  v.upper <-0.6017763*exp((0.8286080/k)*(1/288.15 - 1/(data2$Temp + 273.15))) / 
    (1 + exp((3.1868722/k)*(1/(30.9827367 + 273.15) - 1/(data2$Temp + 273.15))))
  
  v.1<-mean(v.rate)
  v.2<-mean(v.lower)
  v.3<-mean(v.upper)
  v.4<-mean(data2$Temp)
  v.rate.0 <- c(v.rate.0, v.1 )
  v.upper.0 <- c(v.upper.0, v.2 )
  v.lower.0 <- c(v.lower.0, v.3 )
  v.Temp.0 <- c(v.Temp.0, v.4 )
  v.6<-auc(x=seq(1,length(v.rate),1),y=v.rate)/length(v.rate)
  v.int6.0 <- c(v.int6.0,v.6) 
  
  data.F24$Temp<-data2$Temp  
  
}

tab.prms.rate.Y.1  <- data.frame(v.rate.0, v.upper.0, v.lower.0,v.Temp.0,v.int6.0)
data.F24<- read_excel("C:....../Supplementary_data/data/F24.xlsx")

v.rate.0 <- NULL
v.upper.0 <- NULL
v.lower.0 <- NULL
v.Temp.0 <- NULL
v.int6.0<-NULL

for (i in c(1:116)){
  data2 <- subset(data.F24)
  data2$Temp<-data2$Temp -0.1
  
  v.rate<- 0.3584294*exp((0.5810565/k)*(1/288.15 - 1/(data2$Temp + 273.15))) / 
    (1 + exp((2.5680691/k)*(1/(29.2655841 + 273.15) - 1/(data2$Temp + 273.15))))
  v.lower <-  0.4996351  *exp((0.4164426/k)*(1/288.15 - 1/(data2$Temp + 273.15))) / 
    (1 + exp((2.1743233/k)*(1/(26.8258663 + 273.15) - 1/(data2$Temp + 273.15))))
  v.upper <-0.6017763*exp((0.8286080/k)*(1/288.15 - 1/(data2$Temp + 273.15))) / 
    (1 + exp((3.1868722/k)*(1/(30.9827367 + 273.15) - 1/(data2$Temp + 273.15))))
  
  
  v.1<-mean(v.rate)
  v.2<-mean(v.lower)
  v.3<-mean(v.upper)
  v.4<-mean(data2$Temp)
  v.rate.0 <- c(v.rate.0, v.1 )
  v.upper.0 <- c(v.upper.0, v.2 )
  v.lower.0 <- c(v.lower.0, v.3 )
  v.Temp.0 <- c(v.Temp.0, v.4 )
  v.6<-auc(x=seq(1,length(v.rate),1),y=v.rate)/length(v.rate)
  v.int6.0 <- c(v.int6.0,v.6) 
  
  data.F24$Temp<-data2$Temp  
}

tab.prms.rate.Y.2  <- data.frame(v.rate.0, v.upper.0, v.lower.0,v.Temp.0,v.int6.0)
data.Y<- rbind(tab.prms.rate.Y.1,tab.prms.rate.Y.2)
####################C#######
data.F24<- read_excel("C:....../Supplementary_data/data/F24.xlsx")
v.rate.0 <- NULL
v.upper.0 <- NULL
v.lower.0 <- NULL
v.Temp.0 <- NULL
v.int6.0<-NULL

for (i in c(1:116)){
  data2 <- subset(data.F24)
  data2$Temp<-data2$Temp +0.1
  
  v.rate<-  0.2835495*exp((0.5840116/k)*(1/288.15 - 1/(data2$Temp + 273.15))) / 
    (1 + exp((4.1664022/k)*(1/(29.9199098 + 273.15) - 1/(data2$Temp + 273.15))))
  v.lower <-  0.3943706  *exp((0.4513939/k)*(1/288.15 - 1/(data2$Temp + 273.15))) / 
    (1 + exp((3.4050425 /k)*(1/(28.9254233 + 273.15) - 1/(data2$Temp+ 273.15))))
  v.upper <- 0.4448847*exp((0.7358031/k)*(1/288.15 - 1/(data2$Temp + 273.15))) / 
    (1 + exp((5.3859577/k)*(1/(30.7901614 + 273.15) - 1/(data2$Temp + 273.15))))
  
  v.1<-mean(v.rate)
  v.2<-mean(v.lower)
  v.3<-mean(v.upper)
  v.4<-mean(data2$Temp)
  v.rate.0 <- c(v.rate.0, v.1 )
  v.upper.0 <- c(v.upper.0, v.2 )
  v.lower.0 <- c(v.lower.0, v.3 )
  v.Temp.0 <- c(v.Temp.0, v.4 )
  v.6<-auc(x=seq(1,length(v.rate),1),y=v.rate)/length(v.rate)
  v.int6.0 <- c(v.int6.0,v.6) 
  
  data.F24$Temp<-data2$Temp  
  
}

tab.prms.rate.C.1  <- data.frame(v.rate.0, v.upper.0, v.lower.0,v.Temp.0,v.int6.0)
data.F24<- read_excel("C:....../Supplementary_data/data/F24.xlsx")
v.rate.0 <- NULL
v.upper.0 <- NULL
v.lower.0 <- NULL
v.Temp.0 <- NULL
v.int6.0<-NULL


for (i in c(1:116)){
  data2 <- subset(data.F24)
  data2$Temp<-data2$Temp -0.1
  
  v.rate<-  0.2835495*exp((0.5840116/k)*(1/288.15 - 1/(data2$Temp + 273.15))) / 
    (1 + exp((4.1664022/k)*(1/(29.9199098 + 273.15) - 1/(data2$Temp + 273.15))))
  v.lower <-  0.3943706  *exp((0.4513939/k)*(1/288.15 - 1/(data2$Temp + 273.15))) / 
    (1 + exp((3.4050425 /k)*(1/(28.9254233 + 273.15) - 1/(data2$Temp+ 273.15))))
  v.upper <- 0.4448847*exp((0.7358031/k)*(1/288.15 - 1/(data2$Temp + 273.15))) / 
    (1 + exp((5.3859577/k)*(1/(30.7901614 + 273.15) - 1/(data2$Temp + 273.15))))
  
  
  v.1<-mean(v.rate)
  v.2<-mean(v.lower)
  v.3<-mean(v.upper)
  v.4<-mean(data2$Temp)
  v.rate.0 <- c(v.rate.0, v.1 )
  v.upper.0 <- c(v.upper.0, v.2 )
  v.lower.0 <- c(v.lower.0, v.3 )
  v.Temp.0 <- c(v.Temp.0, v.4 )
  v.6<-auc(x=seq(1,length(v.rate),1),y=v.rate)/length(v.rate)
  v.int6.0 <- c(v.int6.0,v.6) 
  
  data.F24$Temp<-data2$Temp  
}

tab.prms.rate.C.2  <- data.frame(v.rate.0, v.upper.0, v.lower.0,v.Temp.0,v.int6.0)
data.C<- rbind(tab.prms.rate.C.1,tab.prms.rate.C.2)

##################S###############
data.F24<- read_excel("C:....../Supplementary_data/data/F24.xlsx")
v.rate.0 <- NULL
v.upper.0 <- NULL
v.lower.0 <- NULL
v.Temp.0 <- NULL
v.int6.0<-NULL

for (i in c(1:116)){
  data2 <- subset(data.F24)
  data2$Temp<-data2$Temp +0.1
  
  v.rate<-  0.2463239*exp((0.5609465/k)*(1/288.15 - 1/(data2$Temp + 273.15))) / 
    (1 + exp((3.6185556/k)*(1/(30.1368399 + 273.15) - 1/(data2$Temp + 273.15))))
  v.lower <-  0.3311925 *exp((0.3983526/k)*(1/288.15 - 1/(data2$Temp + 273.15))) / 
    (1 + exp((2.6578737/k)*(1/(27.8653717 + 273.15) - 1/(data2$Temp + 273.15))))
  v.upper <-0.3937396*exp((0.8431755/k)*(1/288.15 - 1/(data2$Temp + 273.15))) / 
    (1 + exp((4.9434129/k)*(1/(31.2798532 + 273.15) - 1/(data2$Temp + 273.15))))
  
  v.1<-mean(v.rate)
  v.2<-mean(v.lower)
  v.3<-mean(v.upper)
  v.4<-mean(data2$Temp)
  v.rate.0 <- c(v.rate.0, v.1 )
  v.upper.0 <- c(v.upper.0, v.2 )
  v.lower.0 <- c(v.lower.0, v.3 )
  v.Temp.0 <- c(v.Temp.0, v.4 )
  
  v.6<-auc(x=seq(1,length(v.rate),1),y=v.rate)/length(v.rate)
  v.int6.0 <- c(v.int6.0,v.6) 
  
  data.F24$Temp<-data2$Temp  
  
}
data.F24<- read_excel("C:....../Supplementary_data/data/F24.xlsx")
tab.prms.rate.S.1  <- data.frame(v.rate.0, v.upper.0, v.lower.0,v.Temp.0,v.int6.0)


v.rate.0 <- NULL
v.upper.0 <- NULL
v.lower.0 <- NULL
v.Temp.0 <- NULL
v.int6.0<-NULL


for (i in c(1:116)){
  data2 <- subset(data.F24)
  data2$Temp<-data2$Temp -0.1
  
  v.rate<-  0.2463239*exp((0.5609465/k)*(1/288.15 - 1/(data2$Temp + 273.15))) / 
    (1 + exp((3.6185556/k)*(1/(30.1368399 + 273.15) - 1/(data2$Temp + 273.15))))
  v.lower <-  0.3311925 *exp((0.3983526/k)*(1/288.15 - 1/(data2$Temp + 273.15))) / 
    (1 + exp((2.6578737/k)*(1/(27.8653717 + 273.15) - 1/(data2$Temp + 273.15))))
  v.upper <-0.3937396*exp((0.8431755/k)*(1/288.15 - 1/(data2$Temp + 273.15))) / 
    (1 + exp((4.9434129/k)*(1/(31.2798532 + 273.15) - 1/(data2$Temp + 273.15))))
  
  
  v.1<-mean(v.rate)
  v.2<-mean(v.lower)
  v.3<-mean(v.upper)
  v.4<-mean(data2$Temp)
  v.rate.0 <- c(v.rate.0, v.1 )
  v.upper.0 <- c(v.upper.0, v.2 )
  v.lower.0 <- c(v.lower.0, v.3 )
  v.Temp.0 <- c(v.Temp.0, v.4 )
  v.6<-auc(x=seq(1,length(v.rate),1),y=v.rate)/length(v.rate)
  v.int6.0 <- c(v.int6.0,v.6) 
  
  data.F24$Temp<-data2$Temp  
}

tab.prms.rate.S.2  <- data.frame(v.rate.0, v.upper.0, v.lower.0,v.Temp.0,v.int6.0)
data.S<- rbind(tab.prms.rate.S.1,tab.prms.rate.S.2)

###############
data.Y$treatement<-'Y'
data.C$treatement<-'C'
data.S$treatement<-'S'
data.all.24<-rbind(data.Y,data.C,data.S)
############
v.rate.0 <- NULL
v.upper.0 <- NULL
v.lower.0 <- NULL
v.Temp.0 <- NULL
v.int6.0<-NULL

data.F48<- read_excel("C:....../Supplementary_data/data/F48.xlsx")
for (i in c(1:116)){
  data2 <- subset(data.F48)
  data2$Temp<-data2$Temp +0.1
  
  v.rate<- 0.3584294*exp((0.5810565/k)*(1/288.15 - 1/(data2$Temp + 273.15))) / 
    (1 + exp((2.5680691/k)*(1/(29.2655841 + 273.15) - 1/(data2$Temp + 273.15))))
  v.lower <-  0.4996351  *exp((0.4164426/k)*(1/288.15 - 1/(data2$Temp + 273.15))) / 
    (1 + exp((2.1743233/k)*(1/(26.8258663 + 273.15) - 1/(data2$Temp + 273.15))))
  v.upper <-0.6017763*exp((0.8286080/k)*(1/288.15 - 1/(data2$Temp + 273.15))) / 
    (1 + exp((3.1868722/k)*(1/(30.9827367 + 273.15) - 1/(data2$Temp + 273.15))))
  
  v.1<-mean(v.rate)
  v.2<-mean(v.lower)
  v.3<-mean(v.upper)
  v.4<-mean(data2$Temp)
  v.rate.0 <- c(v.rate.0, v.1 )
  v.upper.0 <- c(v.upper.0, v.2 )
  v.lower.0 <- c(v.lower.0, v.3 )
  v.Temp.0 <- c(v.Temp.0, v.4 )
  v.6<-auc(x=seq(1,length(v.rate),1),y=v.rate)/length(v.rate)
  v.int6.0 <- c(v.int6.0,v.6) 
  
  data.F48$Temp<-data2$Temp  
  
}

tab.prms.rate.Y.1  <- data.frame(v.rate.0, v.upper.0, v.lower.0,v.Temp.0,v.int6.0)
data.F48<- read_excel("C:....../Supplementary_data/data/F48.xlsx")

v.rate.0 <- NULL
v.upper.0 <- NULL
v.lower.0 <- NULL
v.Temp.0 <- NULL
v.int6.0<-NULL

for (i in c(1:116)){
  data2 <- subset(data.F48)
  data2$Temp<-data2$Temp -0.1
  
  v.rate<- 0.3584294*exp((0.5810565/k)*(1/288.15 - 1/(data2$Temp + 273.15))) / 
    (1 + exp((2.5680691/k)*(1/(29.2655841 + 273.15) - 1/(data2$Temp + 273.15))))
  v.lower <-  0.4996351  *exp((0.4164426/k)*(1/288.15 - 1/(data2$Temp + 273.15))) / 
    (1 + exp((2.1743233/k)*(1/(26.8258663 + 273.15) - 1/(data2$Temp + 273.15))))
  v.upper <-0.6017763*exp((0.8286080/k)*(1/288.15 - 1/(data2$Temp + 273.15))) / 
    (1 + exp((3.1868722/k)*(1/(30.9827367 + 273.15) - 1/(data2$Temp + 273.15))))
  
  
  v.1<-mean(v.rate)
  v.2<-mean(v.lower)
  v.3<-mean(v.upper)
  v.4<-mean(data2$Temp)
  v.rate.0 <- c(v.rate.0, v.1 )
  v.upper.0 <- c(v.upper.0, v.2 )
  v.lower.0 <- c(v.lower.0, v.3 )
  v.Temp.0 <- c(v.Temp.0, v.4 )
  v.6<-auc(x=seq(1,length(v.rate),1),y=v.rate)/length(v.rate)
  v.int6.0 <- c(v.int6.0,v.6) 
  
  data.F48$Temp<-data2$Temp  
}

tab.prms.rate.Y.2  <- data.frame(v.rate.0, v.upper.0, v.lower.0,v.Temp.0,v.int6.0)
data.Y<- rbind(tab.prms.rate.Y.1,tab.prms.rate.Y.2)
####################C#######
data.F48<- read_excel("C:....../Supplementary_data/data/F48.xlsx")
v.rate.0 <- NULL
v.upper.0 <- NULL
v.lower.0 <- NULL
v.Temp.0 <- NULL
v.int6.0<-NULL

for (i in c(1:116)){
  data2 <- subset(data.F48)
  data2$Temp<-data2$Temp +0.1
  
  v.rate<-  0.2835495*exp((0.5840116/k)*(1/288.15 - 1/(data2$Temp + 273.15))) / 
    (1 + exp((4.1664022/k)*(1/(29.9199098 + 273.15) - 1/(data2$Temp + 273.15))))
  v.lower <-  0.3943706  *exp((0.4513939/k)*(1/288.15 - 1/(data2$Temp + 273.15))) / 
    (1 + exp((3.4050425 /k)*(1/(28.9254233 + 273.15) - 1/(data2$Temp+ 273.15))))
  v.upper <- 0.4448847*exp((0.7358031/k)*(1/288.15 - 1/(data2$Temp + 273.15))) / 
    (1 + exp((5.3859577/k)*(1/(30.7901614 + 273.15) - 1/(data2$Temp + 273.15))))
  
  v.1<-mean(v.rate)
  v.2<-mean(v.lower)
  v.3<-mean(v.upper)
  v.4<-mean(data2$Temp)
  v.rate.0 <- c(v.rate.0, v.1 )
  v.upper.0 <- c(v.upper.0, v.2 )
  v.lower.0 <- c(v.lower.0, v.3 )
  v.Temp.0 <- c(v.Temp.0, v.4 )
  v.6<-auc(x=seq(1,length(v.rate),1),y=v.rate)/length(v.rate)
  v.int6.0 <- c(v.int6.0,v.6) 
  
  data.F48$Temp<-data2$Temp  
  
}

tab.prms.rate.C.1  <- data.frame(v.rate.0, v.upper.0, v.lower.0,v.Temp.0,v.int6.0)
data.F48<- read_excel("C:....../Supplementary_data/data/F48.xlsx")
v.rate.0 <- NULL
v.upper.0 <- NULL
v.lower.0 <- NULL
v.Temp.0 <- NULL
v.int6.0<-NULL


for (i in c(1:116)){
  data2 <- subset(data.F48)
  data2$Temp<-data2$Temp -0.1
  
  v.rate<-  0.2835495*exp((0.5840116/k)*(1/288.15 - 1/(data2$Temp + 273.15))) / 
    (1 + exp((4.1664022/k)*(1/(29.9199098 + 273.15) - 1/(data2$Temp + 273.15))))
  v.lower <-  0.3943706  *exp((0.4513939/k)*(1/288.15 - 1/(data2$Temp + 273.15))) / 
    (1 + exp((3.4050425 /k)*(1/(28.9254233 + 273.15) - 1/(data2$Temp+ 273.15))))
  v.upper <- 0.4448847*exp((0.7358031/k)*(1/288.15 - 1/(data2$Temp + 273.15))) / 
    (1 + exp((5.3859577/k)*(1/(30.7901614 + 273.15) - 1/(data2$Temp + 273.15))))
  
  
  v.1<-mean(v.rate)
  v.2<-mean(v.lower)
  v.3<-mean(v.upper)
  v.4<-mean(data2$Temp)
  v.rate.0 <- c(v.rate.0, v.1 )
  v.upper.0 <- c(v.upper.0, v.2 )
  v.lower.0 <- c(v.lower.0, v.3 )
  v.Temp.0 <- c(v.Temp.0, v.4 )
  v.6<-auc(x=seq(1,length(v.rate),1),y=v.rate)/length(v.rate)
  v.int6.0 <- c(v.int6.0,v.6) 
  
  data.F48$Temp<-data2$Temp  
}

tab.prms.rate.C.2  <- data.frame(v.rate.0, v.upper.0, v.lower.0,v.Temp.0,v.int6.0)
data.C<- rbind(tab.prms.rate.C.1,tab.prms.rate.C.2)

##################S###############
data.F48<- read_excel("C:....../Supplementary_data/data/F48.xlsx")
v.rate.0 <- NULL
v.upper.0 <- NULL
v.lower.0 <- NULL
v.Temp.0 <- NULL
v.int6.0<-NULL

for (i in c(1:116)){
  data2 <- subset(data.F48)
  data2$Temp<-data2$Temp +0.1
  
  v.rate<-  0.2463239*exp((0.5609465/k)*(1/288.15 - 1/(data2$Temp + 273.15))) / 
    (1 + exp((3.6185556/k)*(1/(30.1368399 + 273.15) - 1/(data2$Temp + 273.15))))
  v.lower <-  0.3311925 *exp((0.3983526/k)*(1/288.15 - 1/(data2$Temp + 273.15))) / 
    (1 + exp((2.6578737/k)*(1/(27.8653717 + 273.15) - 1/(data2$Temp + 273.15))))
  v.upper <-0.3937396*exp((0.8431755/k)*(1/288.15 - 1/(data2$Temp + 273.15))) / 
    (1 + exp((4.9434129/k)*(1/(31.2798532 + 273.15) - 1/(data2$Temp + 273.15))))
  
  v.1<-mean(v.rate)
  v.2<-mean(v.lower)
  v.3<-mean(v.upper)
  v.4<-mean(data2$Temp)
  v.rate.0 <- c(v.rate.0, v.1 )
  v.upper.0 <- c(v.upper.0, v.2 )
  v.lower.0 <- c(v.lower.0, v.3 )
  v.Temp.0 <- c(v.Temp.0, v.4 )
  v.6<-auc(x=seq(1,length(v.rate),1),y=v.rate)/length(v.rate)
  v.int6.0 <- c(v.int6.0,v.6) 
  
  data.F48$Temp<-data2$Temp  
  
}
data.F48<- read_excel("C:....../Supplementary_data/data/F48.xlsx")
tab.prms.rate.S.1  <- data.frame(v.rate.0, v.upper.0, v.lower.0,v.Temp.0,v.int6.0)


v.rate.0 <- NULL
v.upper.0 <- NULL
v.lower.0 <- NULL
v.Temp.0 <- NULL
v.int6.0<-NULL


for (i in c(1:116)){
  data2 <- subset(data.F48)
  data2$Temp<-data2$Temp -0.1
  
  v.rate<-  0.2463239*exp((0.5609465/k)*(1/288.15 - 1/(data2$Temp + 273.15))) / 
    (1 + exp((3.6185556/k)*(1/(30.1368399 + 273.15) - 1/(data2$Temp + 273.15))))
  v.lower <-  0.3311925 *exp((0.3983526/k)*(1/288.15 - 1/(data2$Temp + 273.15))) / 
    (1 + exp((2.6578737/k)*(1/(27.8653717 + 273.15) - 1/(data2$Temp + 273.15))))
  v.upper <-0.3937396*exp((0.8431755/k)*(1/288.15 - 1/(data2$Temp + 273.15))) / 
    (1 + exp((4.9434129/k)*(1/(31.2798532 + 273.15) - 1/(data2$Temp + 273.15))))
  
  
  v.1<-mean(v.rate)
  v.2<-mean(v.lower)
  v.3<-mean(v.upper)
  v.4<-mean(data2$Temp)
  v.rate.0 <- c(v.rate.0, v.1 )
  v.upper.0 <- c(v.upper.0, v.2 )
  v.lower.0 <- c(v.lower.0, v.3 )
  v.Temp.0 <- c(v.Temp.0, v.4 )
  v.6<-auc(x=seq(1,length(v.rate),1),y=v.rate)/length(v.rate)
  v.int6.0 <- c(v.int6.0,v.6) 
  
  data.F48$Temp<-data2$Temp  
}

tab.prms.rate.S.2  <- data.frame(v.rate.0, v.upper.0, v.lower.0,v.Temp.0,v.int6.0)
data.S<- rbind(tab.prms.rate.S.1,tab.prms.rate.S.2)

###############
data.Y$treatement<-'Y'
data.C$treatement<-'C'
data.S$treatement<-'S'
data.all.48<-rbind(data.Y,data.C,data.S)

ggplot() +
  geom_line(aes(temp, .fitted, col =treatement), d_preds,size=1.2,show.legend = FALSE ) +
  geom_line(aes(v.Temp.0,v.int6.0),col='goldenrod2',data.all,size=1,show.legend = FALSE)+
  geom_line(aes(v.Temp.0,v.int6.0),col='turquoise4',data.all.24,size=1,show.legend = FALSE)+
  geom_line(aes(v.Temp.0,v.int6.0),col='chartreuse4',data.all.48,size=1,show.legend = FALSE)+
  scale_color_manual(breaks = c("Y", "C", "S","F12","F24","F48"),values=c("Black","Black", "Black" ,"goldenrod2","turquoise4","chartreuse4"),
                     labels=c("Cryptomonas sp.","Chlamydomonas sp","Synechococcus sp","Period 12h","Period 24h","Period 48h"))+
  theme_classic()+ ylim(0.25,0.65)+ xlim(15,35) +theme(axis.text.x = element_text(size=10),axis.text.y = element_text(size=10),axis.title.x = element_text(size=12),axis.title.y = element_text(size=12))+
  labs(x = 'Temperature (ºC)',
       y = 'Growth Rate (d-1)',title='',col='')+
  
  facet_grid(~fct_inorder(treatement))
```
