## Supplemental Tables for "Diet quality alters the consequences of gradual phenotypic plasticity for individual performance"

**Supplementary data**

**Table S1:** Observed temperature mean and variance of the different periodicity treatments

| | Period of fluctuation | Max temperature | Min temperature | Mean temperat ure | Variance |
|---|---|---|---|---|---|
| **F12** | 12 hours | 27.97 | 19.47 | 23.53 | 10.08 |
| **F24** | 24 hours | 27.37 | 18.96 | 22.67 | 11.90 |
| **F48** | 48 hours | 28.01 | 19.56 | 22.68 | 12.19 |

**Table S2: Student test bootstrapping** of the differences between the observed growth rate $\overline{P(T,\Phi)}$ and the mean temperature performance $P(\bar{T},\Phi^{*})$ of *Daphnia magna* under different diets (*Cryptomonas sp*., *Chlamydomonas reinhardtii*., *Synechococcus sp*.) periodicity treatment combinations.

| | Thermal period | df | t | p-value | sign. |
|---|---|---|---|---|---|
| **Cryptomonas sp** | 12h | 10 | 13.68 | <0.001 | *** |
| | 24h | 11 | 16.49 | <0.001 | *** |
| | 48h | 11 | 15.66 | <0.001 | *** |
| **Chlamydomonas reinhardtii** | 12h | 11 | 4.32 | <0.05 | ** |
| | 24h | 11 | 4.95 | <0.001 | *** |
| | 48h | 11 | 6.83 | <0.001 | *** |
| **Synechococcus sp** | 12h | 10 | 2.36 | <0.05 | * |
| | 24h | 11 | -0.01 | 0.99 | ns |
| | 48h | 10 | -0.15 | 0.88 | ns |

**Table S3: Tukey HSD post-hoc comparison** of the observed growth rate $\overline{P(T,\Phi)}$ achieved in the different periodicity treatments.

| | Thermal period 1 | Thermal period 2 | df | f | pvalue | pvalue signif |
|---|---|---|---|---|---|---|
| **Cryptomonas sp** | 12h | 24h | 96 | 0.2444 | 1 | ns |
| | 12h | 48h | 96 | -0.0799 | 1 | ns |
| | 24h | 48h | 96 | -0.331 | 1 | ns |
| **Chlamydomonas reinhardtii** | 12h | 24h | 96 | 3.22 | <0.05 | * |
| | 12h | 48h | 96 | 6.32 | <0.001 | *** |
| | 24h | 48h | 96 | 3.09 | <0.05 | * |
| **Synechcoccus sp** | 12h | 24h | 96 | 1.07 | 0.862 | ns |
| | 12h | 48h | 96 | 0.874 | 1 | ns |
| | 24h | 48h | 96 | -0.177 | 1 | ns |

**Table S4: Tukey HSD post-hoc comparison** of the observed growth rate $\overline{P(T,\Phi)}$ achieved in the different dietary treatments. *Cryptomonas sp* (***Cryp***), *Chlamydomonas reinhardtii* (***Chl***), *Synechococcus sp* (***Syn***).

| | Diet 1 | Diet 2 | df | f | Pvalue adj | pvalue signif |
|---|---|---|---|---|---|---|
| | Chl | Syn | 96 | 8.56 | <0.001 | **** |
| **12h** | Chl | Cryp | 96 | -2.77 | <0.05 | * |
| | Syn | Cryp | 96 | -11.1 | <0.001 | **** |
| | Chl | Syn | 96 | 6.62 | <0.001 | **** |
| **24h** | Chl | Cryp | 96 | -5.81 | <0.001 | **** |
| | Syn | Cryp | 96 | -12.4 | <0.001 | **** |
| | Chl | Syn | 96 | 3.27 | <0.05 | ** |
| **48h** | Chl | Cryp | 96 | -9.23 | <0.001 | **** |
| | Syn | Cryp | 96 | -12.3 | <0.001 | **** |
